# Rapid and automated design of two-component protein nanomaterials using ProteinMPNN

**DOI:** 10.1101/2023.08.04.551935

**Authors:** Robbert J. de Haas, Natalie Brunette, Alex Goodson, Justas Dauparas, Sue Y. Yi, Erin C. Yang, Quinton Dowling, Hannah Nguyen, Alex Kang, Asim K. Bera, Banumathi Sankaran, Renko de Vries, David Baker, Neil P. King

## Abstract

The design of novel protein-protein interfaces using physics-based design methods such as Rosetta requires substantial computational resources and manual refinement by expert structural biologists. A new generation of deep learning methods promises to simplify protein-protein interface design and enable its application to a wide variety of problems by researchers from various scientific disciplines. Here we test the ability of a deep learning method for protein sequence design, ProteinMPNN, to design two-component tetrahedral protein nanomaterials and benchmark its performance against Rosetta. ProteinMPNN had a similar success rate to Rosetta, yielding 13 new experimentally confirmed assemblies, but required orders of magnitude less computation and no manual refinement. The interfaces designed by ProteinMPNN were substantially more polar than those designed by Rosetta, which facilitated *in vitro* assembly of the designed nanomaterials from independently purified components. Crystal structures of several of the assemblies confirmed the accuracy of the design method at high resolution. Our results showcase the potential of deep learning-based methods to unlock the widespread application of designed protein-protein interfaces and self-assembling protein nanomaterials in biotechnology.

## Introduction

Deep learning has revolutionized the field of protein design. Typical design paradigms require three fundamental steps: backbone generation, amino acid sequence design, and structure prediction to evaluate the quality of the designed sequences. Deep learning structure prediction methods such as trRosetta^1^, RoseTTAFold^2^, AlphaFold2^3^ and ESMfold^4^ quickly and accurately generate models of proteins and protein complexes from amino acid sequences. Methods for *de novo* backbone generation such as hallucination, inpainting, and diffusion have significantly enhanced robustness and versatility compared to previous approaches, and have been used to design *de novo* protein monomers, homo-oligomers, proteins bearing functional motifs, and protein- and DNA-binding proteins^5–11^. Likewise, deep learning methods for sequence design such as ABACUS-R^12^, proteinGAN^13^, GVP-GNN^14,15^, and ProteinMPNN^16,17^ have demonstrated exceptional performance both *in silico* and in experimentally characterized proteins, especially monomers and homo-oligomers^18^. In the only reported side-by-side comparisons to date, ProteinMPNN substantially outperformed Rosetta in the design of *de novo* protein homo-oligomers^16^ and binders^19^.

In nature, many protein assemblies with sophisticated functions are constructed from multiple distinct protein subunits or oligomers, which has motivated the development of methods for designing such assemblies for biotechnological applications^20–23^. Rosetta has been a powerful tool for achieving this through protein-protein interface design, but the designed interfaces often rely primarily on hydrophobic packing and require significant manual intervention during the design process to eliminate unnecessary mutations^20,24–27^. While hydrophobic packing provides a strong driving force for assembly, it also tends to make the unassembled protein building blocks prone to aggregation, which can complicate their manufacture. By contrast, the interfaces in naturally occurring hierarchically structured protein complexes often include a higher fraction of polar residues, which maximizes assembly fidelity by minimizing off-target aggregation^28–30^. Methods capable of designing custom multi-component protein assemblies with native-like interfaces would promote the development of new protein-based technologies. For example, a licensed protein nanoparticle vaccine for SARS-CoV-2^31^ uses a variant of the computationally designed two-component icosahedral complex I53-50 that was engineered specifically to enable independent purification and *in vitro* assembly of the two building blocks^25,28^, a feature that was critical for commercial-scale manufacturing.

Here we explore the design of multi-component protein assemblies using ProteinMPNN and establish a fully automated design method that generates novel nanomaterials with high efficiency and accuracy. We benchmark its performance against Rosetta-based design and find that ProteinMPNN generates interfaces with a higher fraction of polar residues, which in several cases yields oligomeric building blocks with favorable solution properties.

## Results

To directly compare the two design methods, we used ProteinMPNN to generate new amino acid sequences for 27 tetrahedral protein assemblies that were previously designed using Rosetta^20^. These assemblies comprise four copies each of two distinct trimeric building blocks, arranged on opposing poles of the three-fold axes of tetrahedral point group symmetry (the “T33” architecture; **Fig. 1**). In the original publication^20^, four of the 27 previously designed complexes successfully adopted the target architecture. Since then, negatively stained electron micrographs of one additional complex, T33-23, revealed monodisperse tetrahedral assemblies of the expected size and morphology following purification (**Fig. S1**).

**Figure 1.**
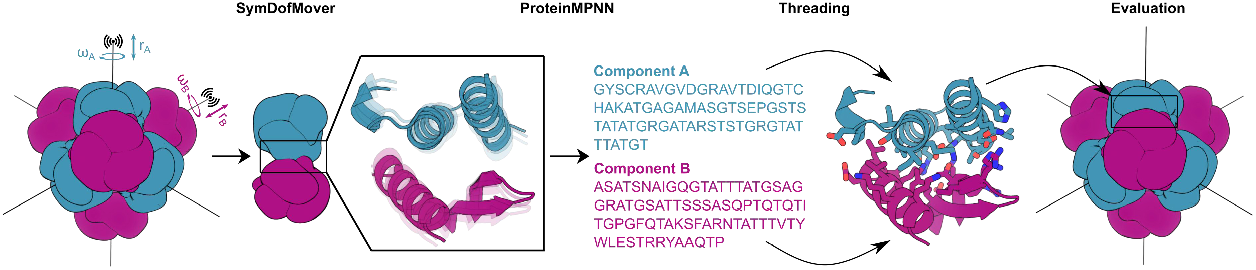
Schematic of two-component tetrahedral nanoparticle design using ProteinMPNN. The rotational (*ω*) and translational (*r*) symmetrical degrees of freedom of a set of previously described T33 docked configurations were sampled using the Rosetta SymDofMover to generate a range of closely related backbones. The backbones of two trimers (one A, one B) were extracted and new amino acid sequences at the A-B interface were generated with ProteinMPNN. The sequences were threaded onto the component backbones using the Rosetta PackRotamerMover to enable evaluation in the context of the full symmetric assembly using Rosetta scoring metrics.

### Nanoparticle design with ProteinMPNN

Our method for designing nanoparticle interfaces using ProteinMPNN is depicted in **Figure 1**. For each of the 27 original T33 designs, we first used the Rosetta SymDofMover to slightly vary the rigid body rotational (*ω*) and translational (*r*) degrees of freedom of each building block. This allowed us to generate 100 docked configurations that were close, but not identical, to the original design. Next, two contacting trimeric components were extracted and ProteinMPNN was used to select the optimal side chain identities for the same sets of interface residues originally considered for design using Rosetta. ProteinMPNN sequence design was rapid, requiring only ∼1 second per sequence, compared to several minutes for Rosetta design. To evaluate the structural features of the designs and enable direct comparison to their Rosetta-designed counterparts, we threaded each ProteinMPNN-designed sequence onto its corresponding dock and evaluated several Rosetta-based interface metrics, including residue counts, clash check, predicted binding energy (ddG), interface surface area, shape complementarity, and the number of buried unsatisfied hydrogen bonding groups. We found that ProteinMPNN and Rosetta yielded designs with roughly similar scores according to these metrics, although the distributions of shape complementarity and predicted binding energy density were slightly better for the Rosetta designs (**Fig. S2**). We selected a maximum of 3 variants of each original design for experimental characterization after ranking the designs by shape complementarity, resulting in a total of 76 designs that passed our filter cut-offs (see Methods) without any manual intervention (**Table S2**). We named these ProteinMPNN-designed nanoparticles by appending a period and a numeric identifier to the name of the original design from which each was derived (e.g., T33-01.1 or T33-25.3).

### Screening and characterization of assembly state

The two components of the 76 designs were encoded as pairs in bicistronic expression plasmids that appended a hexahistidine tag to one of the components. Clarified lysates from 2 mL *E. coli* expression cultures were screened for nanoparticle assembly using polyacrylamide gel electrophoresis (PAGE) under non-denaturing (native) conditions. 24 designs yielded bands that migrated in the range expected of assemblies approaching ∼1 MDa in molecular weight (**Fig. S3**). These 24 potential hits were purified by immobilized metal affinity chromatography (IMAC), re-evaluated by native PAGE, and also analyzed by SDS-PAGE to determine which protein pairs co-eluted, a suggestion of successful assembly (**Fig. S4**). Promising designs were then purified by size-exclusion chromatography (SEC), revealing 13 that eluted as single, symmetric peaks corresponding to ∼1 MDa assemblies (**Fig. 2** and **Fig. S5**). Negatively stained electron micrographs of these 13 designs showed that all of them formed homogeneous assemblies of the expected size and shape (**Fig. 2**). Comparing 2D class averages of each nanoparticle to projections calculated from the computational design models confirmed that all 13 designs assemble to the intended architectures. These results establish that ProteinMPNN can accurately design novel self-assembling protein nanomaterials with a similar success rate to Rosetta—13/76 (17%) vs. 5/27 (18%)—but much more simply and efficiently.

**Figure 2.**
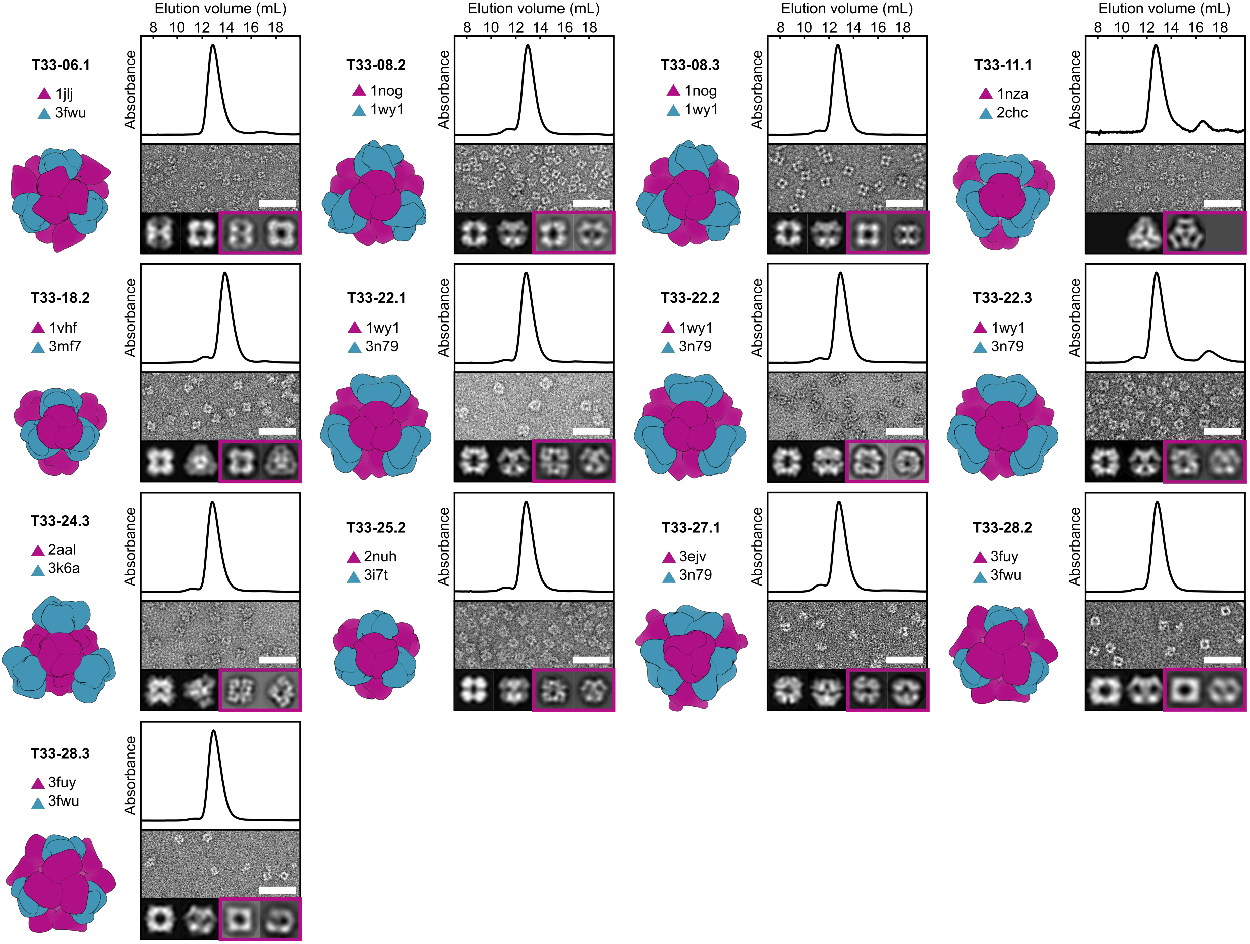
Structural characterization of 13 co-expressed ProteinMPNN-designed tetrahedral nanoparticles. *Left*: Computational design model and the PDB entry from which each of the two trimeric components (A, purple; B, blue) were derived. *Top:* Co-expressed nanoparticles were analyzed by size-exclusion chromatography using a Superdex 200 10/300 column to determine their size and purity. 13 out of the 76 designed nanoparticles eluted at ∼13 mL, consistent with an expected molecular weight (MW) of ∼1 MDa. The absorbance was measured at 230 nm and normalized. *Middle:* Negatively stained electron micrographs of co-expressed tetrahedral nanoparticles. Scale bar: 50 nm. *Bottom left:* Two representative 2D projections calculated from the design model. *Bottom right* (boxed in purple): corresponding experimentally determined 2D class averages. In all cases the 2D class averages closely resemble the 2D projections. In the case of T33-11.1, only a single preferred particle orientation was observed experimentally.

The 13 successful designs were derived from 9 of the 27 distinct two-component protein complexes used as starting points (**Fig. 2**). Two successful designs were obtained from T33-08, two from T33-28, and all three designs based on T33-22 successfully assembled. In these cases, the related design models were highly similar, with backbone root mean square deviations (RMSD) and amino acid sequence identities differing on average by only 0.52 Å over the asymmetric unit (ASU) and 16 distinct amino acid changes out of 45 positions considered for design, respectively. Notably, successful designs were obtained for 8 docked configurations that failed to yield experimentally confirmed assemblies in the original publication. Although ProteinMPNN did not generate successful variants for four of the five previously confirmed T33 nanoparticles (T33-09, T33-15, T33-21, and T33-23), verified assemblies were obtained for 13 of the 27 docked configurations between the two sequence design methods. This high success rate indicates that the simple docking method used— which is available as part of the RPXDock software package^32^—is effective at identifying “designable” docks. One docked configuration yielded successful designs using both Rosetta (T33-28) and ProteinMPNN (T33-28.2 and T33-28.3). Detailed comparisons of the interfaces of these designs are provided below.

### Comparison of ProteinMPNN- and Rosetta-designed interfaces

As an initial comparison of deep learning- and Rosetta-designed interfaces, we visualized the interface residues of each component of the successful ProteinMPNN designs, using color to highlight polar side chains (oxygen atoms colored red, nitrogen atoms colored blue; **Fig. 3a and Fig. S6**). We found that while both sets of interfaces formed well-packed and chemically complementary interactions, the ProteinMPNN-designed interfaces appeared to have more polar residues, especially near the boundary regions of the interface.

**Figure 3.**
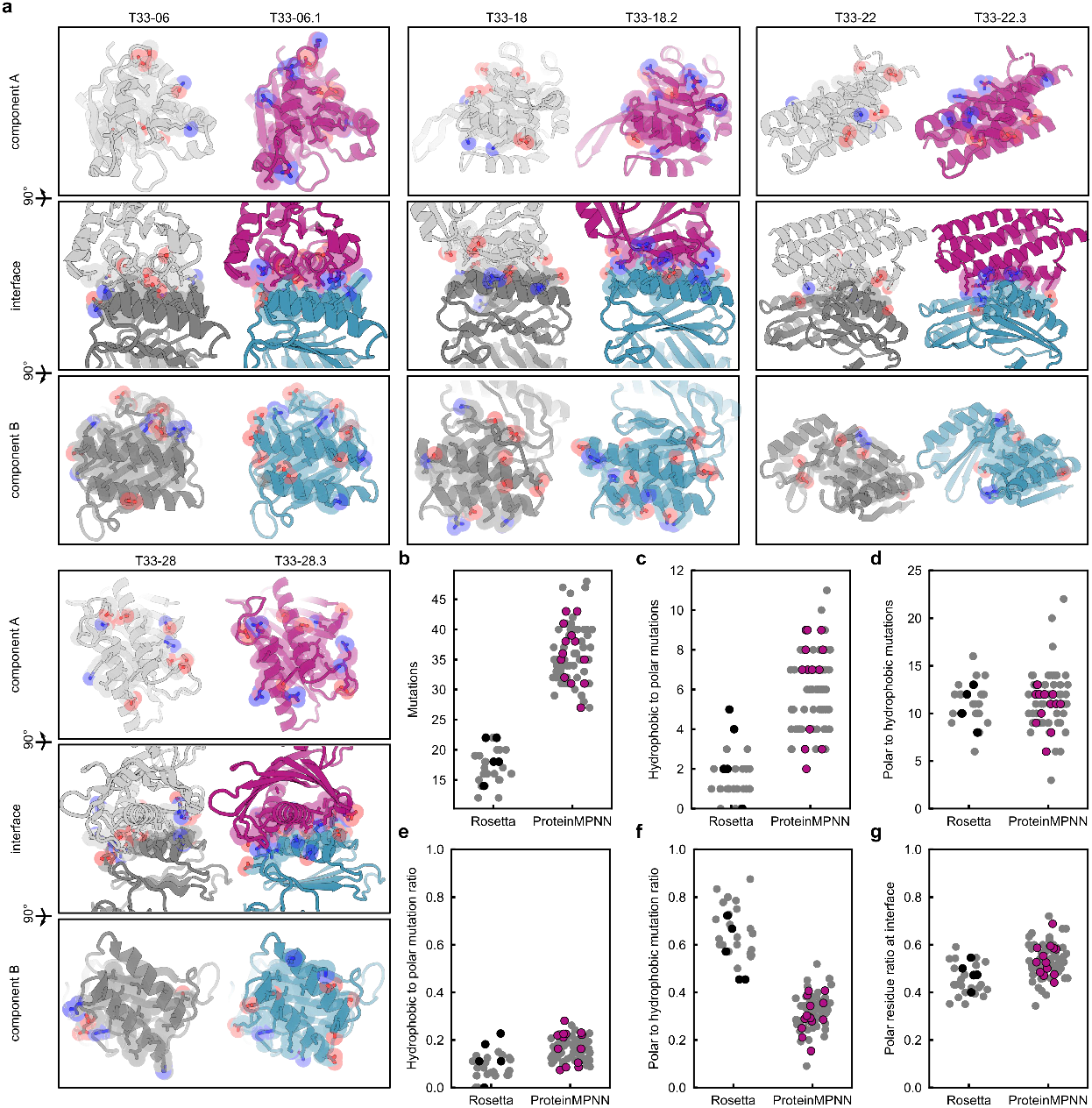
Comparison of Rosetta and ProteinMPNN-Designed interfaces. **(a)** Qualitative comparison of four representative Rosetta-designed interfaces (gray) and ProteinMPNN-designed interfaces (purple/blue). All interface residues are displayed as sticks, with oxygen and nitrogen atoms colored red and blue to highlight polar residues. **(b-g)** Quantitative analyses between the Rosetta (27) and ProteinMPNN design (76) sets, with (**b**) the number of mutations made, **(c)** the number of hydrophobic-to-polar mutations, **(d)** the number of polar-to-hydrophobic mutations, **(e)** the ratio of hydrophobic-to-polar mutations, **(f)** ratio of polar-to-hydrophobic mutations, and **(g)** the ratio of polar residues at the interface. Designs that failed to assemble into target assemblies are shown in gray, while successful designs are shown in black (5 Rosetta designs) or purple (13 ProteinMPNN designs).

We then quantitatively compared the interfaces using several structural metrics. Many of these were similar between the two sets of designs, as noted previously (**Fig. S2**). We found a major difference in the number of mutations to the input scaffolds ProteinMPNN made compared to Rosetta: given the same interface, ProteinMPNN made approximately twice as many mutations (36 ± 5 vs. 17 ± 3; **Fig. 3b**). ProteinMPNN also tended to make more hydrophobic-to-polar mutations (**Fig. 3c**), but a similar number of polar-to-hydrophobic mutations (**Fig. 3d**). When normalized by the total number of mutations, it became clear that ProteinMPNN changed hydrophobes into polars at a similar rate as Rosetta (**Fig. 3e**), but the likelihood of ProteinMPNN converting a polar side chain into a hydrophobic one was much lower (**Fig. 3f**). Quantification of the overall fraction of polar residues at each interface showed that ProteinMPNN designs on average had a higher fraction of polar side chains (**Fig. 3g**). For instance, T33-18.2 has a predominantly polar (59%) interface with only a few well-packed hydrophobes, compared to 43% in the original T33-18 (**Fig. 3a**). This difference between the two methods is even more remarkable considering that the Rosetta designs were manually refined to remove unnecessary polar-to-hydrophobic mutations: the unmodified Rosetta outputs contained even more hydrophobic side chains. This can be explained because the objective of the Rosetta score function is to minimize the energy of the system, and hydrophobic packing is strongly rewarded^33^. Rosetta does not explicitly consider the higher likelihood of oligomeric component aggregation due to surface-exposed hydrophobes, whereas ProteinMPNN was trained on natural protein-protein interfaces that have evolved to balance binding and solubility.

Although none of the metrics evaluated were able to discriminate successful from unsuccessful designs, we note that the ProteinMPNN interfaces also had higher (i.e., worse) predicted binding energies on average according to the Rosetta score function (**Fig. S2**). This result is unsurprising considering that Rosetta explicitly selected mutations that improved Rosetta energy during design. Although experimental determination of interface binding strength would be required to validate these predictions, it is intriguing to consider that studies from our group and others have indicated that relatively weak interfaces can be advantageous for the assembly of protein complexes constructed from oligomeric building blocks^28–30^. This observation, combined with the more polar nature of the ProteinMPNN-designed components, suggested that they may outperform the Rosetta designs in their ability to assemble *in vitro*.

### *In vitro* assembly of ProteinMPNN-designed tetrahedral nanomaterials

The ability to control the assembly of two-component protein nanomaterials by mixing independently purified components *in vitro* simplifies their manufacture and enables advanced functionalization, such as the generation of mosaic nanoparticle immunogens that elicit broadly protective immune responses^34–37^. To evaluate *in vitro* assembly of our 13 ProteinMPNN-designed nanomaterials, we re-cloned each component individually with an appended hexahistidine tag. We were able to successfully purify both components of six complexes by IMAC and SEC (T33-06.1, T33-08.2, T33-11.1, T33-18.2, T33-22.2, and T33-24.3; **Fig. 2**). We mixed the components of each complex in a 1:1 molar ratio at approximately 25 μM in TRIS-buffered saline (TBS; 25 mM Tris pH 8.0, 150 mM NaCl, and 1 mM TCEP), as depicted in **Figure 4a**. All six nanomaterials assembled efficiently and formed monodisperse particles that in negatively stained micrographs were indistinguishable from those obtained by co-expression in *E. coli* (**Fig. 4b**). Dynamic light scattering (DLS) of the assembled materials also indicated efficient formation of the particles of the expected size, with minimal aggregation. By contrast, T33-15 was the only one of the Rosetta designs that could be assembled *in vitro* from purified components^20^.

**Figure 4.**
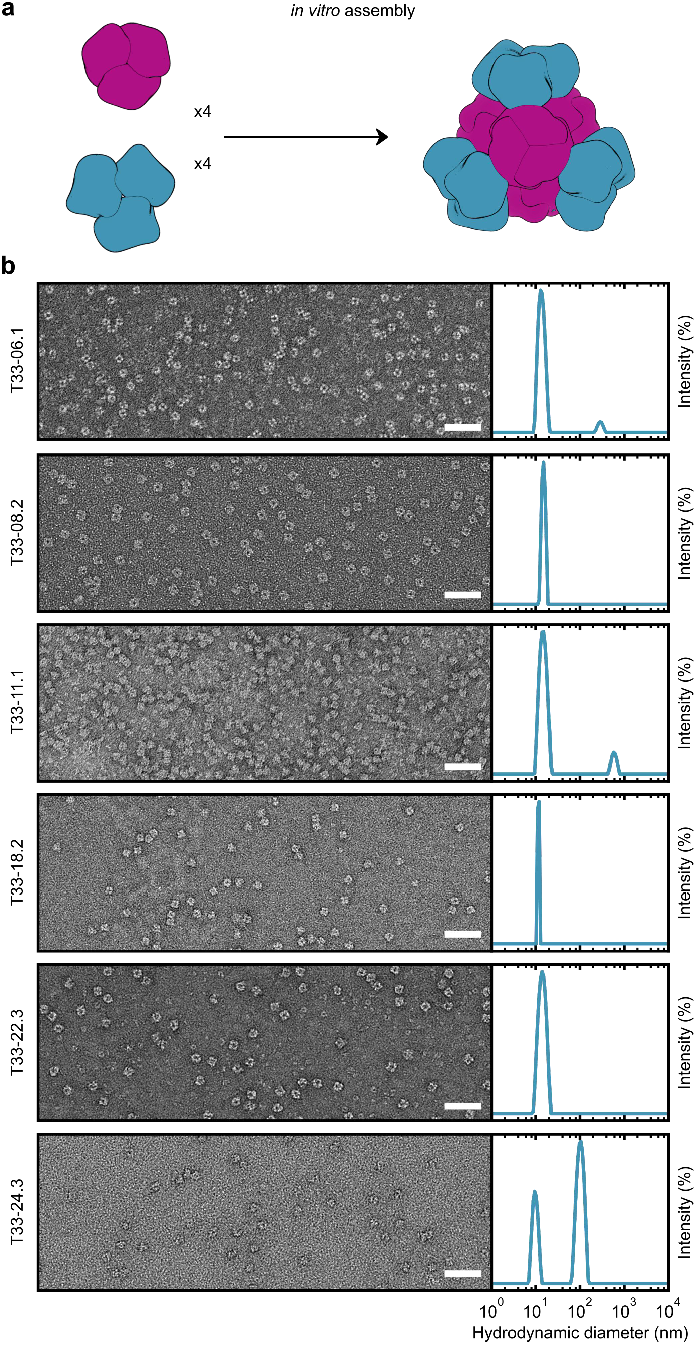
*in vitro* assembly of ProteinMPNN-designed tetrahedral nanoparticles. **(a)** *In vitro* assembly of equimolar components A and B at 25 μM in TBS buffer. **(b)** *Left:* Negatively stained electron micrographs for 6 ProteinMPNN-designed *in vitro* assembled nanoparticles. Scale bar: 50 nm. *Right:* Dynamic light scattering (DLS) of *in vitro* assembled tetrahedral nanoparticles. A scattering peak centered around the expected diameter (∼15 nm) was present for all assembled nanoparticles. For T33-06.1, T33-11.1, and T33-24.3, a second aggregate peak of diameter >100 nm was observed. However, in all instances these peaks represented <1% of the total mass.

**Figure 5.**
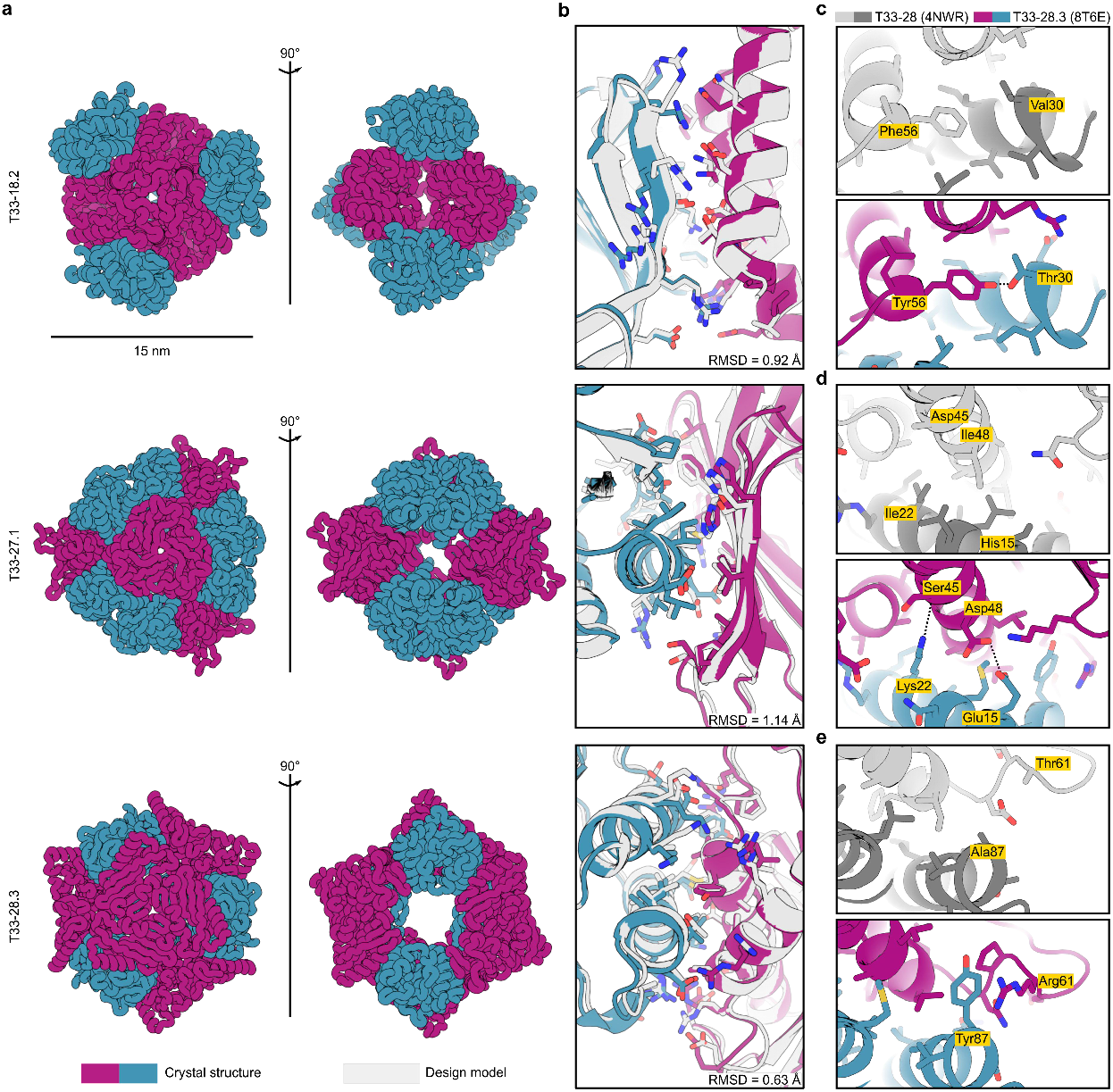
Crystal structures of ProteinMPNN-designed tetrahedral nanoparticles. **(a)** Ribbon displays of crystal structures of three tetrahedral nanoparticles: T33-18.2 (PDB ID: 8T6C), T33-27.1 (PDB ID: 8T6N), and T33-28.3 (PDB ID: 8T6E). **(b)** Atomic interactions at the designed interfaces. Side chains of residues with atoms within 5.5 Å of the opposing subunit are shown as sticks. Light gray color indicates the ProteinMPNN design model and purple-blue colors indicate the crystal structure of components A and B, respectively. In all three instances the crystal structures matched the design models with high accuracy, with backbone Root Mean Square Deviations (RMSD) over the asymmetric unit of: 0.92 Å (T33-18.2), 1.14 Å (T33-27.1), and 0.63 Å (T33-28.3). **(c-e)** Comparison of selected interface interactions in the crystal structures for T33-28 (PDB ID: 4NWR; ref. 20) and T33-28.3 (PDB ID: 8T6E). **(c)** In T33-28.3A the hydrophobic Phe56 is replaced with a Tyr with its hydroxyl group hydrogen bonding with Thr33 on T33-28.3B. **(d)** T33-28.3 features two interface stabilizing polar interactions at the core. The Lys22 amino group forms a hydrogen bond with the backbone carbonyl of Ser45, and Asp48 forms hydrogen bonds with Glu15. In contrast, the core positions for T33-28 (Ile22 and Ile48) featured exclusively hydrophobic packing. **(e)** In a poorly packed region of T33-28, T33-28.3 features a cation-pi interaction between Tyr87 and Arg61. Note: in the crystal structure of T33-28, only the Cβ atom of the His15, Asp45, and Thr61 side chains were resolved.

### High-resolution structure determination

We obtained crystal structures of three of our designs (T33-18.2, T33-27.1, and T33-28.3) to evaluate the accuracy of ProteinMPNN at high resolution (**Fig. 4a and Table S1**). In each case, the crystal structure closely matched the computational design model, with backbone RMSD of 0.92, 1.14, and 0.63 Å over the two-chain asymmetric units of T33-18.2, T33-27.1, and T33-28.3, respectively (**Fig. 4b**). Like other structurally characterized computationally designed assemblies constructed from naturally occurring protein oligomers^20,25,38,39^, these minor deviations largely arise from small rigid body movements of the oligomeric building blocks rather than substantial backbone rearrangements. Although ProteinMPNN does not by default explicitly model side chain configurations during design^16^, we threaded the ProteinMPNN-designed sequences onto each docked configuration using Rosetta to generate full-atom design models. Comparing the side chains of the highest-resolution structure (T33-18.2, with a resolution of 1.92 Å) to its design model revealed that many of the atomic interactions at the designed interface were recapitulated in the crystal structure, both hydrophobic packing interactions as well as hydrogen bonds and electrostatic interactions between polar side chains at the core of the interface (**Fig. 4b**). The configurations of polar side chains around the periphery of the interface deviated more frequently from those predicted in the design model. These data establish that ProteinMPNN can match the high accuracy of Rosetta in the design of novel self-assembling protein nanomaterials.

The crystal structures of T33-28 (ref. 20) and T33-28.3 yielded an opportunity to directly compare experimental structures of interfaces designed by Rosetta and ProteinMPNN. The designs had almost identical overall configurations, with a backbone RMSD of only 0.72 Å over the ASU. However, their interfaces differed by 33 mutations, of which 11 were at core positions. For example, a key core interaction in both assemblies features an aromatic side chain at position 56 in T33-28A that is packed into a complementary hydrophobic groove on the other subunit. ProteinMPNN placed a Tyr in this position, with its hydroxyl forming a hydrogen bond with Thr30 on T33-28B, instead of the more hydrophobic Phe selected by Rosetta (**Fig. 4c**). Two additional hydrophobic-to-polar mutations form hydrogen bonds at the core of the T33-28.3 interface: Ile22Lys in T33-28.3B and Ile48Asp in T33-28.3A (**Fig. 4d**). Several more polar residues at the boundary regions of T33-28.3 allowed the formation of more than 10 favorable hydrogen and ionic interactions, compared to zero polar interactions across the T33-28 interface (**Fig. S7**). Most of these interactions were accurately recapitulated in the T33-28.3 crystal structure, although at the periphery some deviations of polar rotamer conformations were observed due to interaction with water molecules. Finally, a poorly packed region on the periphery of the T33-28 interface was redesigned by ProteinMPNN to feature a tyrosine residue from T33-28.3B (Tyr87) that is involved in several favorable packing interactions across the interface, including a cation-pi interaction with Arg61 from T33-28.3A that was observed in the crystal structure (**Fig. 4e**). The latter interaction was present in all three ProteinMPNN designs, while cation-pi interactions are rarely designed by Rosetta. In addition to highlighting the different types of interactions designed by each algorithm, these comparisons between the Rosetta- and ProteinMPNN-designed T33-28 assemblies establish how highly divergent designed interface sequences can drive the assembly of nearly geometrically identical self-assembling protein materials.

## Discussion

Computationally designed self-assembling proteins are a promising technology platform that has begun to yield commercial products. Two-component nanoparticles similar to those reported here have been used to encapsulate and deliver molecular cargoes^40–42^, to scaffold proteins for structural characterization by cryo-electron microscopy^43–45^, and to display antigens in repetitive arrays that, when used as vaccines, elicit robust and in some cases broadly protective immune responses^22,34,35,46–49^. The recent licensure of a protein nanoparticle vaccine for SARS-CoV-2 established computationally designed self-assembling proteins as a commercial technology^31,50^. Nevertheless, the technology is still relatively new, and the assemblies designed to date are relatively simple. By contrast, the remarkably sophisticated self-assembling proteins observed in nature—and the highly specialized functions they perform—hint at the technological potential of designed protein assemblies. Continued methods development will help realize this potential by making possible the design of synthetic protein assemblies that rival the structural and functional complexity of the molecular machines of the cell.

Many recent developments in computational methods for modeling and designing proteins have focused on machine learning^4,51–57^. ProteinMPNN^16^, based on the earlier graph-based Message-Passing Neural Network (MPNN) architecture of Ingraham et al. (ref. 17), has proven to be a robust and versatile tool for fixed-backbone sequence design: ProteinMPNN can be used as a sequence design module in a wide variety of protein redesign and *de novo* design tasks. As we have shown, this includes the design of novel self-assembling protein nanomaterials, where ProteinMPNN outperformed Rosetta, the previous state-of-the-art sequence design method, in a head-to-head comparison. Although the two methods generated experimentally confirmed assemblies with similar success rates, the speed and simplicity of ProteinMPNN are considerable advantages. In particular, its modest computational resource requirements and its elimination of the need for intensive manual review by structural biologists should make it accessible to a wider set of researchers than Rosetta, facilitating the application of protein design as a solution to a wider variety of challenges in biology and beyond.

We found an additional advantage of ProteinMPNN in its ability to design protein-protein interfaces with a higher proportion of polar residues than Rosetta. This directly translated to improved biophysical properties in the oligomeric components of our two-component protein nanomaterials. Specifically, the lower tendency of ProteinMPNN-designed components to aggregate resulted in a higher fraction of materials that could be assembled *in vitro* from individually purified components. *In vitro* assembly has been used in the manufacturing processes for several designed protein nanoparticle vaccines in clinical trials^35,49^ (NCT05664334 & NCT04750343), including a mosaic nanoparticle vaccine for influenza that co-displays four distinct hemagglutinin antigens on the same nanoparticle immunogen^34^ (NCT04896086). Optimizing this property of designed protein nanomaterials is therefore of potential commercial relevance. Furthermore, as new computational methods enable the design of increasingly complex protein nanomaterials—such as those that break symmetry or comprise several different components^58–60^—hierarchical *in vitro* assembly will become even more preferred as a method of construction, mirroring the reticular synthesis of complex metal-organic frameworks and DNA nanotechnology objects^61–64^. Our results suggest that the ability of ProteinMPNN to design components of protein nanomaterials with favorable solution properties will likely speed the development of the next generation of protein-based technologies.

## Materials and Methods

### Computational design

The symmetrical degrees of freedom of the T33 nanoparticle docks were sampled with Δ0.5 Å translation (*r*) and Δ1° rotation (*ω*) using the Rosetta SymDofMover, and 100 unique configurations per dock were generated. Because for some docks the interface spans multiple subunits of the homotrimers, the trimers of both components sharing a single nanoparticle interface were isolated, and the identical interface residues to King et al. (ref. ^20^) were identified. The interface sequences were optimized using ProteinMPNN, with all other residues kept fixed. A total of 16 sequences per sampled configuration (1,600 sequences per dock) were generated with ProteinMPNN using sampling temperatures of 0.2 and 0.3, and backbone noise of 0.05. Based on the loss score, the top 50% of the sequences were selected and threaded back onto the sampled asymmetric units using Rosetta resfiles, and nanoparticle design models were generated using the SymDofMover. All residues were repacked using the Rosetta PackRotamerMover (the packer) with tetrahedral symmetry. Finally, a detailed evaluation of the designed interfaces was performed by filtering interfaces based on the following metrics: number of glycines = 0, number of methionines < 5, number of aromatics < 5, number of clashes < 3, shape complementarity > 0.5, predicted interface strength < 0 Rosetta Energy Units (REU), and solvent accessible surface area buried at the designed interface > 1000 Å^2^. For each of the 27 docks, the 3 highest shape complementarity designs were selected, with the exceptions of T33-15—where only 1 design passed the filter metrics—and T33-12, T33-16, and T33-29—where only 2 designs passed the filter metrics. This gave a total of 76 designs for the ProteinMPNN-designed set.

### Bicistronic expression, lysate screening, and purification

Synthetic genes for designed proteins optimized for *E. coli* expression were purchased from Genscript ligated into the pET-29b(+) vector using the NdeI and XhoI restriction sites. A second ribosome binding site was inserted between the open reading frames of the two components of the bicistronic nanoparticle designs (AGAAGGAGATATCAT) such that the two proteins would be co-expressed. Only one of the components with the most accessible C-terminus carried a hexahistidine tag to facilitate co-purification. Plasmids were cloned into BL21 (DE3) *E. coli* competent cells (New England Biolabs). Transformants were grown in 2 mL lysogeny broth (LB; 10 g Tryptone, 5 g Yeast Extract, 10 g NaCl) cultures in 96 deep-well plates at 37 °C for 2 h, induced with 1 mM IPTG, and then continued to shake at 37 °C for ∼18 h. The cells were harvested and lysed by sonication using a plate sonicator (Qsonica) for 2 min in pulses of 20 sec on, 40 sec off in 25 mM Tris pH 8.0, 300 mM NaCl, 30 mM imidazole, 1 mM PMSF, 0.05% (w/v) DNAse I. Lysates were clarified by centrifugation for 15 min at 4,000 *g* in a swinging bucket rotor. Supernatants were applied to native PAGE gels and run at 100 V for ∼4 h on ice. Potential nanoparticle hits had bands at ∼1 MDa after staining with GelCode (Thermo Fisher Scientific).

The potential hits were expressed in 1 L LB cultures in 2 L baffled shake flasks. The cells were harvested and lysed by sonication using a microplate horn system (Qsonica) for 10 minutes with 10 s pulses at 80% amplitude in 25 mM Tris pH 8.0, 300 mM NaCl, 30 mM imidazole, 1 mM PMSF, 0.05% (w/v) DNAse I. Lysates were clarified by centrifugation at 24,000 *g* for 30 min and applied to a 2 mL column bed of Ni-NTA resin (Qiagen) for purification by IMAC. The resin was washed with 50 mL buffer 25 mM Tris pH 8.0, 300 mM NaCl, 30 mM imidazole. The protein of interest was eluted using 6 mL of 25 mM Tris pH 8.0, 300 mM NaCl, 300 mM imidazole. Directly after elution, 1 mM dithiothreitol (DTT; Sigma Aldrich) was added to eluates. Eluates were evaluated for co-elution by SDS-PAGE and assembly by native PAGE. IMAC eluates that contained nanoparticle bands on native PAGE were concentrated in 100 kDa MWCO centrifugal filters (Amicon), sterile filtered (0.22 μm), and applied to a Superdex 200 Increase 10/300 (Cytiva) in 25 mM Tris pH 8.0, 150 mM NaCl, 1 mM DTT. Peaks eluting at ∼13 mL strongly indicate nanoparticle formation, and were fractionated and analyzed by SDS-PAGE to confirm the presence of both components.

### Individual component expression and purification

Synthetic genes for individual components, each with a hexahistidine purification tag, were optimized for *E. coli* expression and purchased from Genscript ligated into the pET-29b(+) vector at the NdeI and XhoI restriction sites. The proteins were expressed in BL21(DE3) (New England Biolabs) in LB grown in 2 L baffled shake flasks. Cells were grown at 37 °C to an OD_600_ ∼ 0.6, and then induced with 1 mM IPTG. Expression temperature was reduced to 18 °C and the cells were shaken for ∼18 h. The cells were harvested and lysed by sonication using a Qsonica Q500 for 10 min with 10 s pulses at 80% amplitude in 25 mM Tris pH 8.0, 300 mM NaCl, 30 mM imidazole, 1 mM PMSF, 0.05% (w/v) DNAse I. Lysates were clarified by centrifugation at 24,000 *g* for 30 min and applied to a 5 mL column bed of Ni-NTA resin (Qiagen) for purification by IMAC. Resin was pre-washed with 25 mM Tris pH 8.0, 300 mM NaCl, 30 mM imidazole. The protein of interest was eluted using 25 mM Tris pH 8.0, 300 mM NaCl, 300 mM imidazole. DTT was added to eluates to a final concentration of 1 mM. Eluates were pooled, concentrated in 10K MWCO centrifugal filters (Pall), sterile filtered (0.22 μm) and applied to either a Superdex 200 Increase 10/300 (Cytiva), or HiLoad 26/600 Superdex 200 pg SEC column (Cytiva) using 25 mM Tris pH 8.0, 300 mM NaCl, 1 mM TCEP buffer.

### *In vitro* assembly

Total protein concentration of purified individual nanoparticle components was determined by measuring absorbance at 280 nm using a UV/vis spectrophotometer (Agilent Cary 3500 Multicell) and calculated extinction coefficients^65^. The assembly steps were performed at room temperature with addition in the following order: component A, followed by additional buffer as needed to achieve desired final concentration, and finally component B (in 25 mM Tris pH 8.0, 150 mM NaCl), with a molar ratio of A:B of 1:1. T33-08.2, T33-11.1, and T33-18.2 components were incubated at room temperature for at least 1 h in order to drive a more complete assembly. These assemblies were applied to a Superose 6 Increase 10/300 GL column (Cytiva) for purification by SEC using 25 mM Tris pH 8.0, 300 mM NaCl, 1 mM TCEP running buffer. Assembled nanoparticles were sterile filtered (0.22 μm) immediately prior to column application and following pooling of fractions. T33-06.1, T33-22.3, and T33-24.3 assembly reactions were incubated at room temperature for 18 h, then sterile filtered (0.22 μm) and analyzed directly, without subsequent SEC.

### Negative stain electron microscopy collection and processing

Tetrahedral nanoparticles were first diluted to 100 μg/mL in water prior to application of 3 μL of sample onto freshly glow-discharged 400-mesh copper grids (Ted Pella). Sample was incubated on the grid for 30 s before excess liquid blotted away with filter paper (Whatman). 3 μL of 2% w/v uranyl formate (UF) stain was applied to the grid and immediately blotted away before an additional 3 μL of UF stain was applied. Stain was blotted off by filter paper, and a final 3 μL of UF stain was applied and allowed to incubate for ∼30 s. Finally, the stain was blotted away and the grids were allowed to dry for 3 min. Prepared grids were imaged using EPU 2.0 on a 120 kV Talos L120C transmission electron microscope (Thermo Fisher Scientific) at 57,000× magnification with a BM-Ceta camera. Data processing was done in CryoSPARC^66^, starting with CTF correction, particle picking and extraction. Two or three rounds of 2D classification were done.

### Crystallographic data collection and structure determination

All crystallization experiments were conducted using the sitting drop vapor diffusion method. Crystallization trials were set up in 200 nL drops using the 96-well plate format at 20 °C. Crystallization plates were set up using a Mosquito LCP from SPT Labtech, then imaged using UVEX microscopes and UVEX PS-256 from JAN Scientific. Diffraction quality crystals formed in 0.9 M NPS 0.1 M Tris-BICINE pH 8.5, 30 % v/v of Glycerol and PEG 4000 for T33-18.2; 0.12 M Ethylene glycol, 0.1 M Tris-BICINE pH 8.5, 30 % v/v of Glycerol and PEG 4000 for T33-27.1; and 1.26 M Sodium phosphate, 0.14 M Potassium phosphate for T33-28.3.

Diffraction data was collected at the Advanced Light Source (ALS) HHMI beamline 8.2.1/8.2.2 and 5.0.1. at 1Å wavelength. X-ray intensities and data reduction were evaluated and integrated using XDS^67^ and merged/scaled using Pointless/Aimless in the CCP4 program suite^67,68^. Starting phases were obtained by molecular replacement using Phaser^69^ using the designed model for the structures. Following molecular replacement, the models were improved using phenix.autobuild^70^; efforts were made to reduce model bias by setting rebuild-in-place to false, and using simulated annealing and prime-and-switch phasing. Structures were refined in Phenix^70^. Model building was performed using COOT^71^. The final model was evaluated using MolProbity^72^. Data collection and refinement statistics are recorded in **Table S1**. Data deposition, atomic coordinates, and structure factors reported in this paper have been deposited in the Protein Data Bank (PDB)^73^, http://www.rcsb.org/, with accession codes **8T6C** (T33-18.2), **8T6N** (T33-27.1), and **8T6E** (T33-28.3).

## Acknowledgements

This work was funded by the Bill & Melinda Gates Foundation (INV-010680 to D.B. and N.P.K.), the National Science Foundation (DMREF 1629214 to D.B. and N.P.K.), the National Institute of Allergy and Infectious Disease (U54AI170856 and 1P01AI167966 to N.P.K.), the Howard Hughes Medical Institute (D.B.), the Audacious Project at the Institute for Protein Design (D.B. and N.P.K.), and the Open Philanthropy Project Improving Protein Design Fund (D.B). This work was also supported financially by the VLAG Graduate School Research Fellowship and Fulbright Visiting Scholar Fellowship to R.J.dH. We thank the Advanced Light Source (ALS) beamline 8.2.1/8.2.2/5.0.1 at Lawrence Berkeley National Laboratory for X-ray crystallography data collection. The Berkeley Center for Structural Biology is supported in part by the National Institutes of Health (NIH), National Institute of General Medical Sciences, and the Howard Hughes Medical Institute. The ALS is supported by the Director, Office of Science, Office of Basic Energy Sciences and US Department of Energy (DOE) (DE-AC02-05CH11231).

## Competing Interests

The authors declare no competing interests.

## Notes

### Competing Interest Statement

The authors have declared no competing interest.

