## Supplementary Information for "Rapid and automated design of two-component protein nanomaterials using ProteinMPNN"

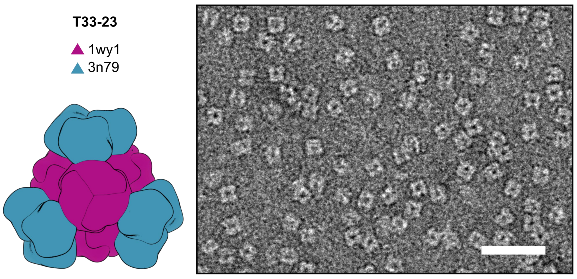

**Figure S1. Negative stain electron microscopy of purified T33-23.** *Left*: Computational design model and the PDB entry from which each of the two trimeric components (A, purple; B, blue) were derived. *Right*: Negatively stained electron micrograph of the co-expressed tetrahedral nanoparticle. Scale bar: 50 nm.

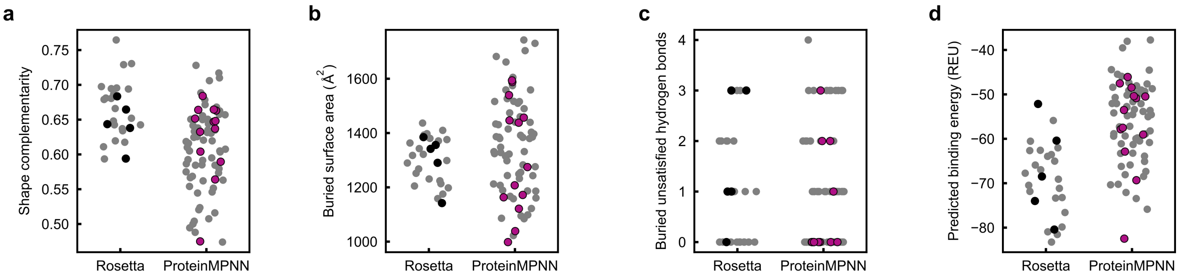

**Figure S2. Detailed scoring metrics of tetrahedral nanoparticle interface design with Rosetta and ProteinMPNN.** Interface scoring metrics plotted: **(a)** shape complementarity, **(b)** buried surface area in Å$^2$, **(c)** buried unsatisfied hydrogen bonds, and **(d)** predicted binding energy (ddG) in Rosetta Energy Units (REU).

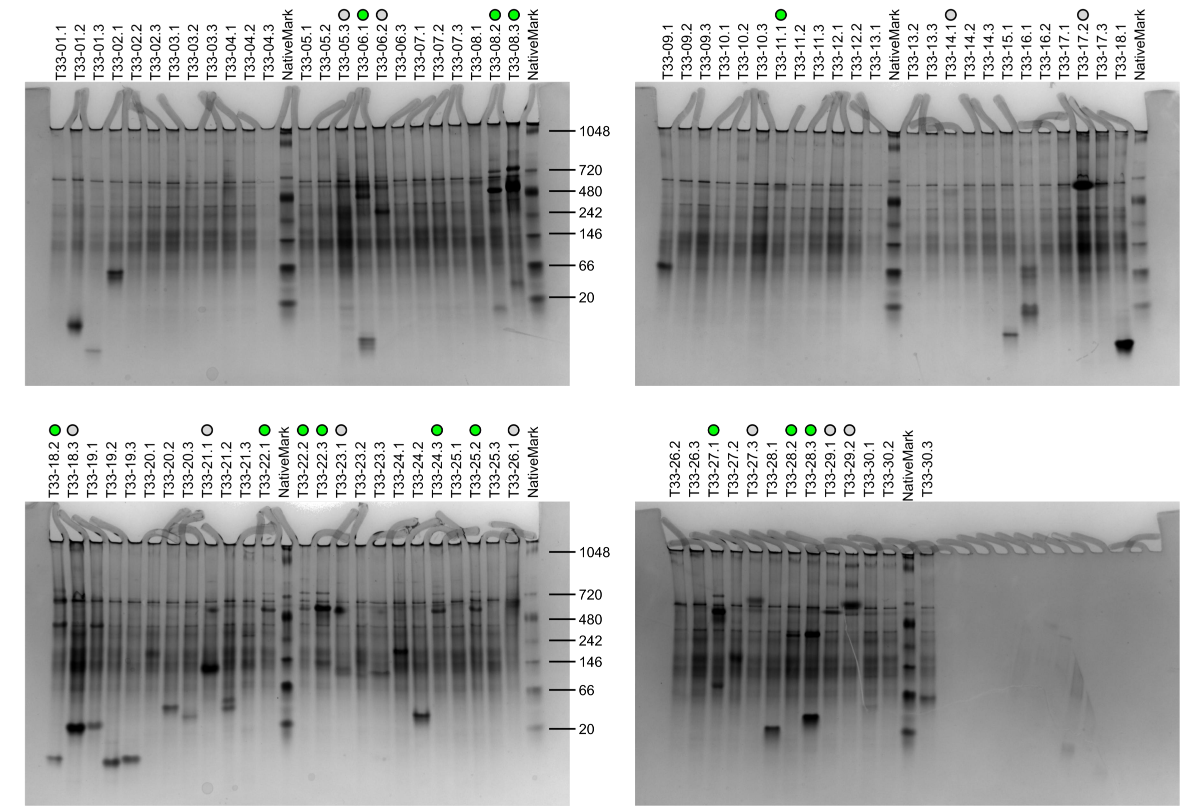

**Figure S3. Lysate screening of bicistronically expressed ProteinMPNN-designed tetrahedral nanoparticles.** Assembly into tetrahedral nanoparticles results in slower migration in native PAGE. 24 Designs yielding bands that migrated in ~0.5-1 MDa range were selected for further characterization. Circles indicate selected designs, green circles indicate designs that were eventually confirmed to assemble into tetrahedral nanoparticles. NativeMark (ThermoFisher Scientific) is loaded as reference with molecular weight in kDa as indicated. Note: apart from the 76 designs representing the 27 experimentally characterized docks described in King et al. (ref. [20]), an additional 9 ProteinMPNN designs were tested based on T33-19, T33-20, and T33-30, which were designed but not experimentally tested in King et al. None of the 9 ProteinMPNN designs exhibited indications of nanoparticle assembly. To ensure fair comparison of docks for which experimental data was available for both the Rosetta and ProteinMPNN design sets, these 9 designs were not further considered.

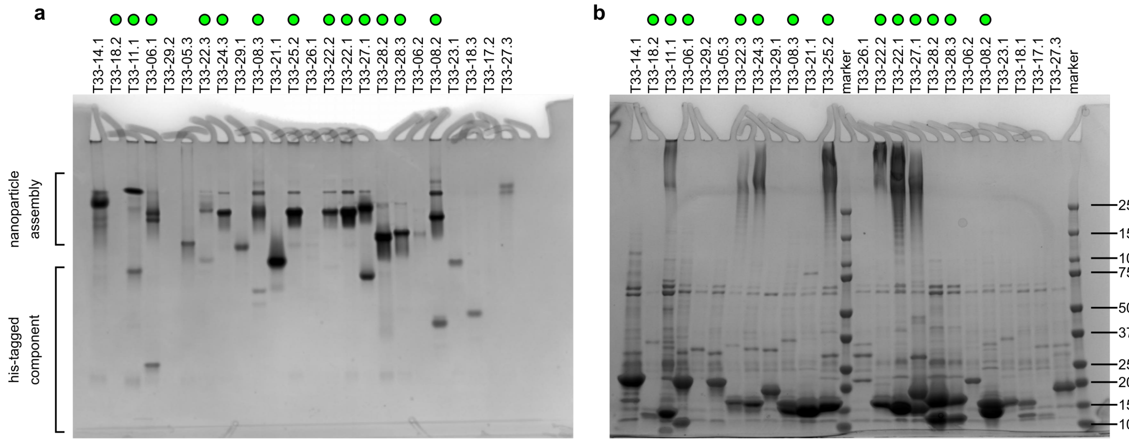

**Figure S4. Co-purification of 24 potential ProteinMPNN-designed tetrahedral nanoparticles. (a)** (Non)-denaturing PAGE of Ni-NTA purified tetrahedral assembly candidates. Assembled nanoparticles migrated slower in the gel compared to individual components. **(b)** Denaturing PAGE of Ni-NTA purified tetrahedral nanoparticle candidates. A and B components with different molecular weights (between ~10 and ~20 kDa) can be observed, indicating co-purification since only one of the two components contained a hexahistidine tag. Protein marker with molecular weights indicated in kDa is shown. Green colored circles indicate designs that were eventually confirmed to assemble into tetrahedral nanoparticles.

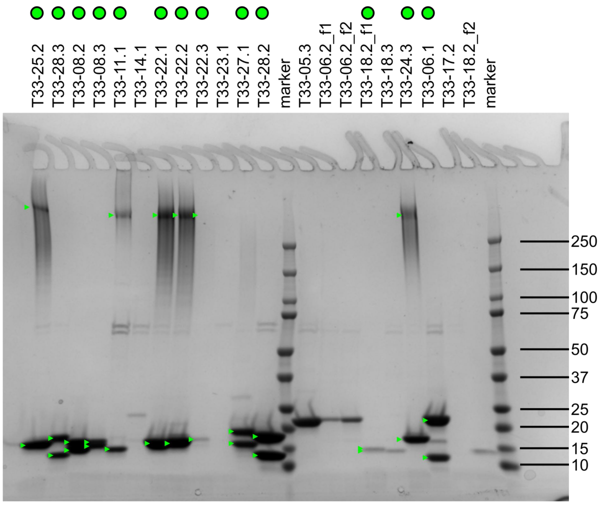

**Figure S5. Denaturing PAGE of isolated size exclusion chromatography nanoparticle peaks.** Nanoparticle peak (~13 mL) of a superdex S200 10/300 Increase column (Cytiva) is isolated. “f1” indicates peak isolated at ~13 mL and “f2” indicates peak isolated at ~14 mL. Marker with molecular weight indicated in kDa in figure. Green triangles indicate the two components. Green colored circles indicate designs that were eventually confirmed to assemble into tetrahedral nanoparticles. Note that for T33-25.2, T33-28.3, T33-11.1 T33-22.1, T33-22.2, T33-22.3 and T33-24.3 one of the two components does not fully migrate into the gel.

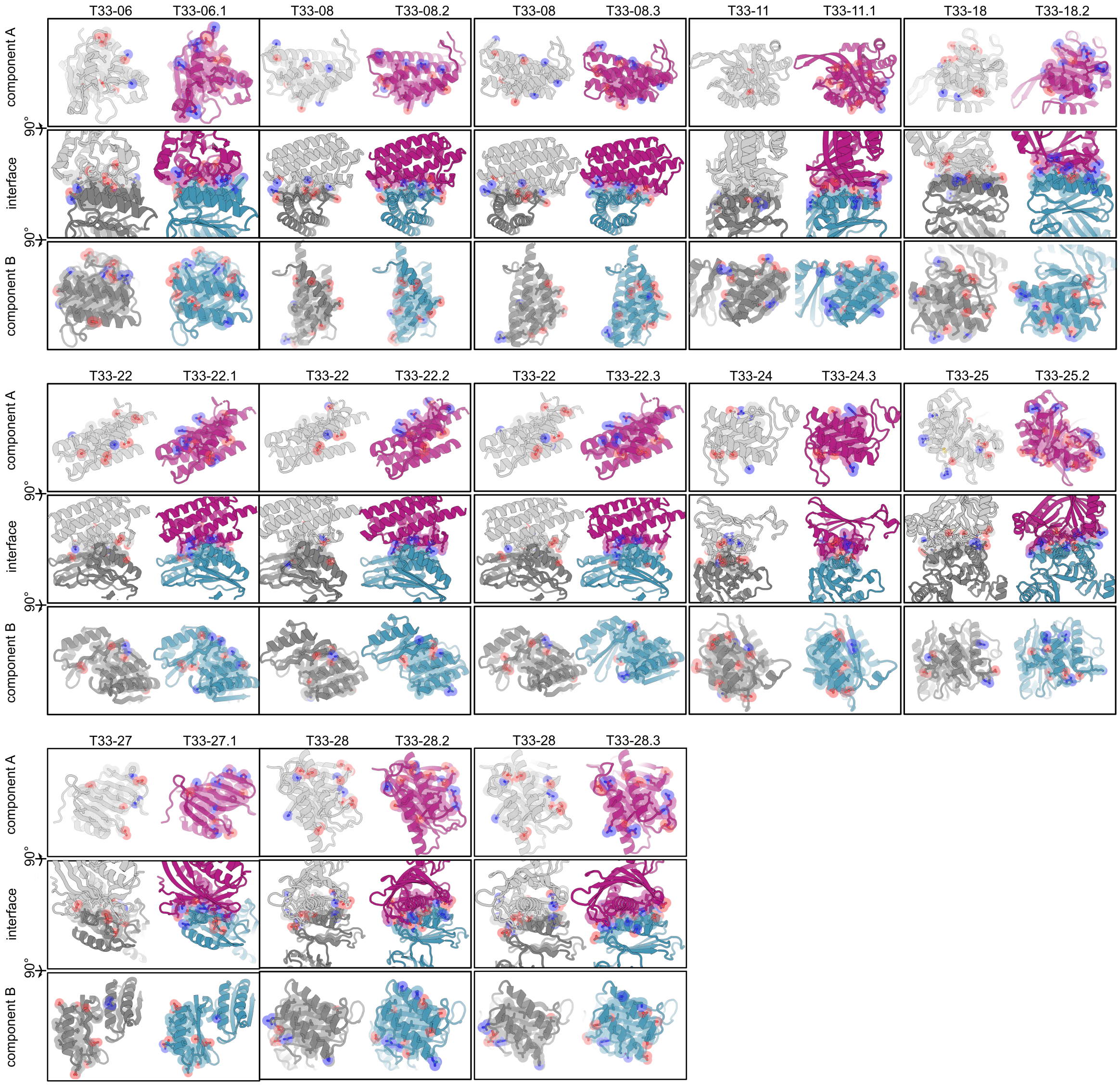

**Figure S6. Comparison of Rosetta-designed interfaces and ProteinMPNN-designed interfaces**. for all 13 confirmed ProteinMPNN-designed tetrahedral nanoparticles. All interface residue side chains within 5.5 Å distance across the interface are displayed as sticks, with oxygen and nitrogen atoms colored red and blue respectively, to highlight polar residues.

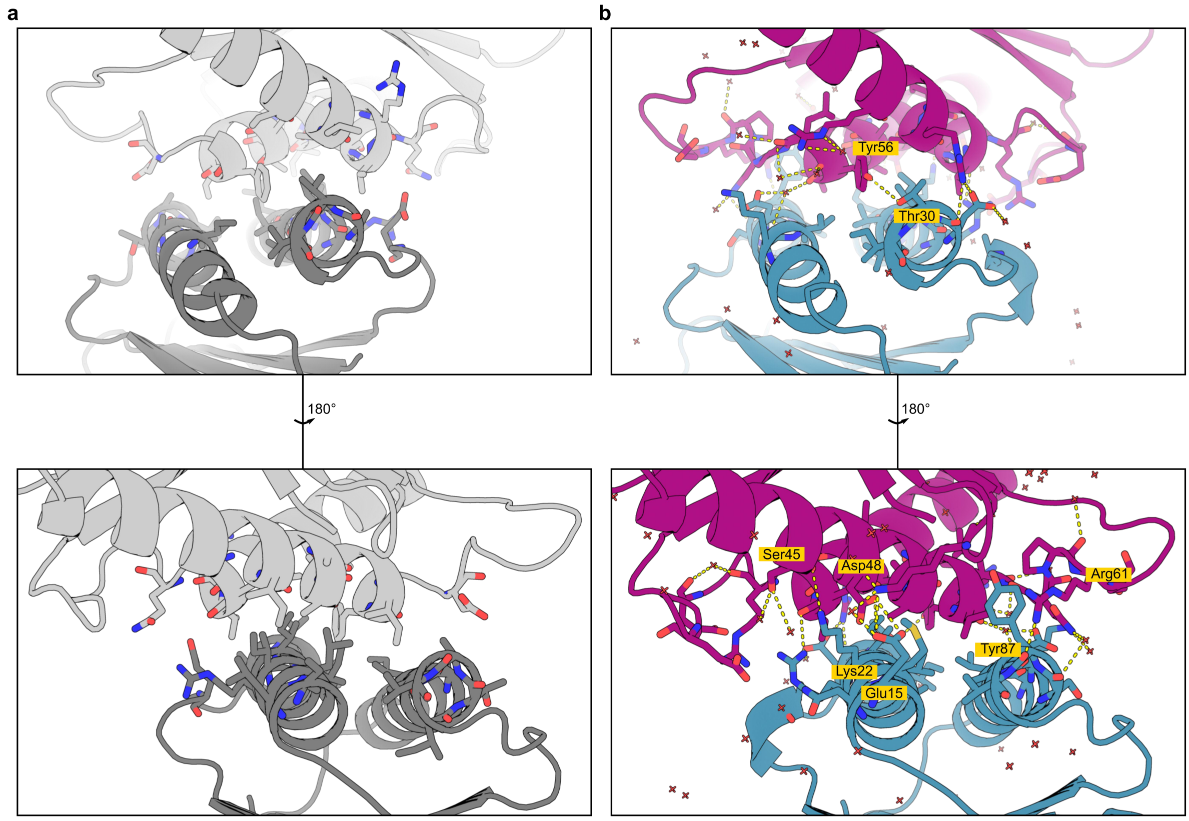

**Figure S7. Interface comparison of crystal structure of T33-28 and T33-28.3.** Two orientations at 180° of the interface of crystal structures T33-28 (PDB ID: 4NWR[20]) **(a)** and T33-28.3 (PDB ID: 8T6E) **(b)** are shown. All residues with heavy atoms at less than 5.5 Å across the interface are visualized as sticks. All possible hydrogen bonding and ionic interactions across the interface and with water molecules at less than 3.5 Å distance were visualized (yellow dotted lines). For the T33-28 crystal structure zero polar interactions were observed, but for T33-28.3 more than 10 interactions across the interface were present. Key polar mutations are highlighted in yellow in T33-28.3. Water molecules are visualized as red plusses.

**Table S1. Data collection and refinement statistics (molecular replacement).**

| | **T33-18.2 (8T6C)** | **T33-27.1 (8T6N)** | **T33-28.3 (8T6E)** |
|---|---|---|---|
| **Data collection** | | | |
| Space group | | | |
| Cell dimensions | | | |
| *a, b, c* (Å) | 138.81,138.81, 129.11 | 176.51, 176.51, 176.5 | 113.74, 114.05, 114.20 |
| **a, b, g (°)** | 90, 90, 120 | 90.00, 90.00, 90.00 | 62.09, 77.86, 89.41 |
| Resolution (Å) | 56.87 - 1.92 (1.96 - 1.92)* | 100 - 3.63 (3.69 - 3.63)* | 48.47 - 2.48 (2.52 - 2.48)* |
| $R_{sym}$ or $R_{merge}$ | 0.085 (0.162) | 0.229 (0.063) | 0.062 (0.630) |
| I / sI | 9.8 (0.1) | 22.69 (1.5) | 6.3 (0.7) |
| Completeness (%) | 100 (100) | 100 (100) | 98.1 )97.5) |
| Redundancy | 6.7 (6.7) | 21.5 (21.4) | 2.4 (2.2) |
| **Refinement** | | | |
| Resolution (Å) | 56.87 - 1.92 (1.95 - 1.92) | 48.96 - 3.63 (3.72 - 3.63) | 48.47 - 2.48 (2.54 - 2.48) |
| No. reflections | 70444 | 20763 | 171172 |
| $R_{work}$ / $R_{free}$ | 0.1977 (0.4685)/ 0.2423 (0.5712) | 0.2142 (0.2654)/ 0.2585 (0.3624) | 0.2053 (0.3513)/ 0.2389 (0.4115) |
| No. atoms | | | |
| Protein | 7211 | 1175 | 25202 |
| Ligand/ion | n/a | n/a | n/a |
| Water | 424 | n/a | 593 |
| B-factors | | | |
| Protein | 38.44 | 91.68 | 55.66 |
| Ligand/ion | n/a | n/a | n/a |
| Water | 40.79 | n/a | 50.57 |
| R.m.s. deviations | | | |
| Bond lengths (Å) | 0.004 | 0.002 | 0.002 |
| Bond angles (°) | 0.600 | 0.432 | 0.460 |

*A single crystal was used for each structure in the data collection. Values in parentheses are for the highest-resolution shell.

**Table S2. Amino acid sequences of novel proteins designed in this study.**

| Design | Component A | Component B | MW$^{componentA}$ (kDa) | MW$^{componentB}$ (kDa) |
|---|---|---|---|---|
| T33-01.1 | MPIFTLNTNIKATDVPSD FLARTSRLVALILNKPGS YVAVHINTDQQLSFGGS TNPAAFGTLMSIGGIEP GLNHAINCALTAELEHEL GIAPERMYIHFVNLNGD DVGWGYGTG | MTEKEKMLAEKWYDANF DQELIERRAKAAICWAL NNTRPSDTGRIKALIDALF GKVTDNVSISIPFDTDYG ENVKLGKNVYVNTNCYF MDGGQITIGDNVFIGPNC GFYTATHPLNFHHRNEG FEKAGPIHIGSNTWFGGH VAVLPGVTIGEGSVIGAG SVVTKDIPPHSLAVGNPC KVVRKIDNDEGSHHHHH H | 12.1 | 21.5 |
| T33-01.2 | MPIFTLNTNIKATDVPSD FLARTSEAVSRILNKPG SYVAVHINTDQQLSFGG STNPAAFGTLMSIGGIEP ENNEKLNVVLTTLLEHE LGIPADRMYIHFVNLNG DDVGWGYGTG | MTEKEKMLAEKWYDANF DQELIARRTRAAILCYILN NTRPSDKELRKLLIDALF RTKTDNVSISIPFDTDYG ENVKLGKNVYVNTNCYF MDGGQITIGDNVFIGPNC GFYTATHPLNFHHRNEG FEKAGPIHIGSNTWFGGH VAVLPGVTIGEGSVIGAG SVVTKDIPPHSLAVGNPC KVVRKIDNDEGSHHHHH H | 12.3 | 21.8 |
| T33-01.3 | MPIFTLNTNIKATDVPSD FLDVTSRLVADLLEKPG SYVAVHINTDQQLSFGG STNPAAFGTLMSIGGIEP GLNAAINAALTAVLEDLL GIRGDRMYIHFVNLNGD DVGWGYGTG | MTEKEKMLAEKWYDANF DQELIAERARAEAICWAL NNTRPSNKGRIRGLIEGL FGKVTDNVSISIPFDTDY GKNVKLGKNVYVNTNCY FMDGGQITIGDNVFIGPN CGFYTATHPLNFHHRNE GFEKAGPIHIGSNTWFG GHVAVLPGVTIGEGSVIG AGSVVTKDIPPHSLAVGN PCKVVRKIDNDEGSHHH HHH | 12.0 | 21.5 |
| T33-02.1 | MHQIRVGVLTVSDSCFR NLAPDRSGEALKLYVQD KDELGGTISAYKIVPDEI EEIKETLIDWCDEKELNL ILTTGGTGFAPRDVTPE ATKEVIEREAPGMALAM LMASLSLSPEGMLSRPV CGIRGKTLIINLPGSLIGS LHCFDAILPALPHAIDLL RDAIVKVKEVHLGSHHH HHH | MPVIQTFVSTPLDHHKRE ELAEVYRRVTREVLGKP EDLVMMTFHDSTPMHFF GSTDPVACVRVEALGGY KENQPSVVTRTVSAAISL ECGIVLERIFVLYFSPLHC GWNGTNP | 19.4 | 12.7 |
| T33-02.2 | MHQIRVGVLTVSDSCFR NLAKDRSGNALKWLVQ | MPVIQTFVSTPLDHHKRE NLAEVYRAVTREILGKPE | 19.5 | 12.8 |

| | | | | |
|---|---|---|---|---|
| | DPDLLGGTISAYKIVPDE IEEIKETLIDWCDEKELN LILTTGGTGFAPRDVTP EATKEVIEREAPGMALA MLMASLEVSPEGMLSR PVCGIRGKTLIINLPGST VGSLRCFLAILPALPHAI DLLRDAIVKVKEVHHGS HHHHHH | DLVMMTFHDSTPMHFFG STDPVACVRVEALGGYR EGEPELVTLIVTYAIEDEC GIVKDRIFVLYFSPLHCG WNGTNE | | |
| T33-02.3 | MHQIRVGVLTVSDSCFR NLAPDWSGNALKRLVQ DPELLGGTISAYKIVPDE IEEIKETLIDWCDEKELN LILTTGGTGFAPRDVTP EATKEVIEREAPGMALA MLMHSLSVSPRGMLSR PVCGIRGKTLIINLPGSQI GSLRCFLAILPALPHAID LLRDAIVKVKEVHHGSH HHHHH | MPVIQTFVSTPLDHHKRE ALAERYRAVTKEILGKPE DLVMMTFHDSTPMHFFG STDPVACVRVEALGGYK EDEPELVTLIVTLSIEEEC GIVKERIFVLYFSPLHCG WNGTNP | 19.5 | 12.8 |
| T33-03.1 | MHQIRVGVLTVSDSCFR NLERPLSGRALELYVQD PKLLGGTISAYKIVPDEIE EIKETLIDWCDEKELNLIL TTGGTGFAPRDVTPEAT KEVIEREAPGMALAMLM GSLNITPLGMLSRPVCGI RGKTLIINLPGSLLGSVR CFKFILPALPHAIDLLRD AIVKVKEVHHGSHHHHH H | MPVIQTFVSTPLDELRRE SLVLVYRLVTEEVLGKPA DLVMMTFHDSTPMHFFG STDPVACVRVEALGGYG PSEPEKVTEIVTRAITEVC GIVADRIFVLYFSPLHCG WNGTNL | 19.6 | 12.5 |
| T33-03.2 | MHQIRVGVLTVSDSCFR NLIPPLSGEALKIFVQDP KLLGGTISAYKIVPDEIEE IKETLIDWCDEKELNLILT TGGTGFAPRDVTPEATK EVIEREAPGMALAMLMG SLNVTPLGMLSRPVCGI RGKTLIINLPGSLLGSLQ CFQFILPALPHAIDLLRD AIVKVKEVHHGSHHHHH H | MPVIQTFVSTPLDELRRQ SLVLTYRIVTEKILGKPAD LVMMTFHDSTPMHFFGS TDPVACVRVEALGGYGP SEPEKVTEIVTRAITEVCG IVADRIFVLYFSPLHCGW NGTNL | 19.5 | 12.5 |
| T33-03.3 | MHQIRVGVLTVSDSCFR NLRRPLSGRALEIFVQD PELLGGTISAYKIVPDEIE EIKETLIDWCDEKELNLIL TTGGTGFAPRDVTPEAT KEVIEREAPGMALAMLM GSLNLTPLGMLSRPVCG IRGKTLIINLPGSLLGSLS CFRFILPALPHAIDLLRD AIVKVKEVHEGSHHHHH H | MPVIQTFVSTPLDELRRE SLCLVYRIVTEEILGKPAD LVMMTFHDSTPMHFFGS TDPVACVRVEALGGYGP SEPERVTRVVTEAITEVC GIVADRIFVLYFSPLHCG WNGTNL | 19.6 | 12.6 |
| T33-04.1 | MHQIRVGVLTVSDSCFR | MPVIQTFVSTPLDEDRRE | 19.6 | 12.5 |

| | | | | |
|---|---|---|---|---|
| | NLRPDLSGKALEILVQD<br>PAFLGGTISAYKIVPDEI<br>EEIKETLIDWCDEKELNL<br>ILTTGGTGFAPRDVTPE<br>ATKEVIEREAPGMALAM<br>LMGSLKITPLGMLSRPV<br>CGIRGKTLIINLPGSLLG<br>SLRCFRFILPALPHAIDLL<br>RDAIVKVKEVHHGSHHH<br>HHH | ALCLVYRIVTEEILGKPAD<br>LVMMTFHDSTPMHFFGS<br>TAPVACVRVEALGGYGP<br>SEPERVTEIVTKAITEVCG<br>IVADRIFVLYFSPLHCGW<br>NGTNL | | |
| T33-04.2 | MHQIRVGVLTVSDSCFR<br>NLRPDLSGRALARYVQ<br>DPRELGGTISAYKIVPDE<br>IEEIKETLIDWCDEKELN<br>LILTTGGTGFAPRDVTP<br>EATKEVIEREAPGMALA<br>MLMGSLSITPLGMLSRP<br>VCGIRGKTLIINLPGSLL<br>GSLECFRFILPALPHAID<br>LLRDAIVKVKEVHHGSH<br>HHHHH | MPVIQTFVSTPLNEARRE<br>ALVLVYRLVTKEILGKPE<br>DLVMMTFHDSTPMHFFG<br>STAPVACVRVEALGGYG<br>PSEPEKVTAVVTEAITEV<br>CGIVADRIFVLYFSPLHC<br>GWNGTNL | 19.6 | 12.4 |
| T33-04.3 | MHQIRVGVLTVSDSCFR<br>NLRPDLSGRALRRYVQ<br>DPAELGGTISAYKIVPDE<br>IEEIKETLIDWCDEKELN<br>LILTTGGTGFAPRDVTP<br>EATKEVIEREAPGMALA<br>MLMGSLKITPLGMLSRP<br>VCGIRGKTLIINLPGSLL<br>GSLRCFRFILPALPHAID<br>LLRDAIVKVKEVHHGSH<br>HHHHH | MPVIQTFVSTPLDEDRRE<br>ALFLVYQLVTEEILGKPA<br>DLVMMTFHDSTPMHFFG<br>STAPVACVRVEALGGYG<br>PSEPEKVTEVVTKAITEV<br>CGIVADRIFVLYFSPLHC<br>GWNGTNL | 19.7 | 12.5 |
| T33-05.1 | MHQIRVGVLTVSDSCFR<br>NLATDWSGEALKLYVSN<br>PKRLGGTISAYKIVPDEI<br>EEIKETLIDWCDEKELNL<br>ILTTGGTGFAPRDVTPE<br>ATKEVIEREAPGMALAM<br>LMGSLAVTPLGMLSRPV<br>CGIRGKTLIINLPGSLSG<br>SLRCFEFILPALPHAIDLL<br>RDAIVKVKEVHEGSHHH<br>HHH | MPVIQTFVSTPLDHHKRE<br>NLAQAYRDVTRRILGKPE<br>DLVMMTFHDSTPMHFFG<br>STDPVACVRVEALGGYG<br>RGEPELVTLVVSLAIEKE<br>CGIVLERIFVLYFSPLHCG<br>WNGINF | 19.5 | 12.7 |
| T33-05.2 | MHQIRVGVLTVSDSCFR<br>NLARDRSGRALRLYVAN<br>PKELGGTISAYKIVPDEI<br>EEIKETLIDWCDEKELNL<br>ILTTGGTGFAPRDVTPE<br>ATKEVIEREAPGMALAM<br>LMGSLAVTPLGMLSRPV<br>CGIRGKTLIINLPGSLRG<br>SLRCFEFILPALPHAIDLL<br>RDAIVKVKEVHHGSHHH<br>HHH | MPVIQTFVSTPLDHHKRE<br>NLAQVYRDVTREILGKPE<br>DLVMMTFHDSTPMHFFG<br>STDPVACVRVEALGGYG<br>KDEPELVNLVVSLAIQEE<br>CGIVLDRIFVLYFSPLHCG<br>WNGINV | 19.6 | 12.7 |

| | | | | |
|---|---|---|---|---|
| T33-05.3 | MHQIRVGVLTVSDSCFRNLAEDWSGRALELYVTNPNELGGTISAYKIVPDEIEEIKETLIDWCDEKELNLILTTGGTGFAPRDVTPEATKEVIEREAPGMALAMLMGSLKVTPLGMLSRPVCGIRGKTLIINLPGSLEGSLRCFRFILPALPHAIDLLRDAIVKVKEVHAGSHHHHHH | MPVIQTFVSTPLDHHKRENLAQVYRDVTRRILGKPEDLVMMTFHDSTPMHFFGSTDPVACVRVEALGGYGREQPELVTLVVSLAIDKECGIVLDRIFVLYFSPLHCGWNGINF | 19.6 | 12.8 |
| T33-06.1 | MHQIRVGVLTVSDSCFRNLRPDLSGRALERYVQDPKLLGGTISAYKIVPDEIEEIKETLIDWCDEKELNLILTTGGTGFAPRDVTPEATKEVIEREAPGMALAMLMGSLNITPLGMLSRPVCGIRGKTLIINLPGSLLGSLRCFDFILPALPHAIDLLRDAIVKVKEVHHGSHHHHH | MPVIQTFVSTPLDEDDRRALSLVYRYATEKILGKPADLVMMTFHDSTPMHFFGSTDPVACVRVEALGGYGPSEPEEVTKLVTAAITEVCGIVADRIFVLYFSPLHCGWNGTNV | 19.6 | 12.4 |
| T33-06.2 | MHQIRVGVLTVSDSCFRNLRPDLSGEALKIFVQDPELLGGTISAYKIVPDEIEEIKETLIDWCDEKELNLILTTGGTGFAPRDVTPEATKEVIEREAPGMALAMLMGSLNRTPLGMLSRPVCGIRGKTLIINLPGSLLGSLRCFRFILPALPHAIDLLRDAIVKVKEVHHGSHHHHHH | MPVIQTFVSTPLDEDDRTALCLVYRIVTERILGKPADLVMMTFHDSTPMHFFGSTDPVACVRVEALGGYGPSEPEEVTRVVTAAITEVCGIVADRIFVLYFSPLHCGWNGTNV | 19.6 | 12.4 |
| T33-06.3 | MHQIRVGVLTVSDSCFRNLRPDLSGEALRRIVQDPALLGGTISAYKIVPDEIEEIKETLIDWCDEKELNLILTTGGTGFAPRDVTPEATKEVIEREAPGMALAMLMGSLNITPLGMLSRPVCGIRGKTLIINLPGSLLGSLACFRFILPALPHAIDLLRDAIVKVKEVHEGSHHHHHH | MPVIQTFVSTPLDELDRASLDLVYRIATERILGKPADLVMMTFHDSTPMHFFGSTDPVACVRVEALGGYGPSEPEEVTRLVTEAITEVCGIVADRIFVLYFSPLHCGWNGTNV | 19.5 | 12.4 |
| T33-07.1 | MALYFMGHMILVYSTFPSEKIAEITGKALLAQRLIACFNAFEIRSGYWWKGEVVQDKEWAAIFKTTEEKEKELYEALRELHPEETPAIFTLKVENILTEYMNWLRESVLGE | MQAIGILELTSIAKGMELGDAMLKSANVDLLVSKTISPGKFLLMLGGDVPAIAKAVIVGIGNAGEMLVDSRVILDIHPSVLPAISGLNSVDKRQAVGIVETWSVAACIKAADAAVKGSNVTLVRVHMAFGIGGKCYMVVAGDVSDVNNAVTVASESAGEKGLLVYRSVIPRPHEAMWRQMVEGGSHHHHHH | 13.0 | 20.0 |

| | | | | |
|---|---|---|---|---|
| T33-07.2 | MALYFMGHMILVYSTFP NELLAEITGKALLAKRLIA CFNAFEIRSGYWWKGEI VQDKEWAAIFKTTEEKE KELYEALREVHVFETPAI FTLKVENILTEYMNWLR ESVLGG | MSQAIGILELTSIAKGMEL GDAMLKSANVDLLVSKTI SPGKFLLMLGGDVLAIAK AVAVGLGAAGEMLVDSR LITNIHPSVLPAISGLNSV DKRQAVGIVETWSVAACI KAADKAVKGSNVTLVRV HMAFGIGGKCYMVVAGD VSDVNNAVTVASESAGE KGLLVYRSVIPRPHEAM WRQMVEGGSHHHHHH | 13.0 | 20 |
| T33-07.3 | MALYFMGHMILVYSTFP NEAQAQIVGKALLAQRLI ACFNAFEIRSGYWWKG EIVQDKEWAAIFKTTEEK EKELYEALRELHVEETP AIFTLKVENILTEYMNWL RESVLGA | MSQAIGILELTSIAKGMEL GDAMLKSANVDLLVSKTI SPGKFLLMLGGDVLAIAK AVSVGLGNAGEMLVDSA LLTNIHPSVLPAISGLNSV DKRQAVGIVETWSVAACI KAADKAVKGSNVTLVRV HMAFGIGGKCYMVVAGD VSDVNNAVTVASESAGE KGLLVYRSVIPRPHEAM WRQMVEGGSHHHHHH | 13.0 | 20 |
| T33-08.1 | MSPVVEVQGTIDELNSFI GYALVLSRWDDIRNDLF RIQNDLFVLGEDVSTGG KGRTVTLEMILDLVKKTE KMLKEIGKIELFVVPGGS VESASLHMARAVSRRLE RRIRAAAELTPINALVEL YAEALSKILFMHALISNK RLNIPEKIL | MREPIIEANGTLDELTSFI GEAKHYVDEEMKGILEEI QNDIYKIMGEIGSKGKIEG IPLSSLNRLVDLIERYEEM VNKSFVLPGGTLESAKLD VCRTIARRALLKVETVLR EFGIGLVAVLYLEVLSELL FLLARVIEIEKNKKGSHH HHHH | 16.8 | 17.2 |
| T33-08.2 | MSPVVEVQGTIDELNSFI GYALVLSRWDDIRNDLF RIQNDLFVLGEDVSTGG KGRTVTLEMIAELVKKS YKMKKEIGKIELFVVPG GSVESASLHMARAVSR RLERRIEAAAKLTEINEL VLLYAQALSRILFMHALI SNKRLNIPEKIY | MNQPIIEANGTLDELTSFI GEAKHYVDEEMKGILEEI QNDIYKIMGEIGSKGKIEG ISDESLVKLLDLIERYEEM VNKSFVLPGGTLESAKLD VCRTIARRATLKVKTVLE EFGIGFNAVLYLEVLSELL FLLARVIEIEKNKEGSHH HHHH | 16.9 | 17.2 |
| T33-08.3 | MSPVVEVQGTIDELNSFI GYALVLSRWDDIRNDLF RIQNDLFVLGEDVSTGG KGRTVTEEMVIELIKRYV KMKKEIGKIELFVVPGG SVESASLHMARAVSRRL ERRIEAAARLTPINELVL AYAQALSRILFMHALISN KRLNIPEKIL | MNQPIIEANGTLDELTSFI GEAKHYVDEEMKGILEEI QNDIYKIMGEIGSKGKIEG ISDESVVKLWDLIERYEE MVNKSFVLPGGTLESAK LDVCRTIARRAYLKVLTV VREFGIGLTAVLYLKLLSE LLFLLARVIEIEKNKEGSH HHHHH | 16.9 | 17.3 |
| T33-09.1 | MEEVVLITVPSAEEAVRI AYTLVEERLAACVNIIPG VVKIYRWQGRVRVASTL LLLVKTTTHAFPKLKERV KALHPYTVPEIVALPIAE | MVRGIRGAITVAADTDEA ILAATIELLREMLRANGIQ SYEELAAVIFTVTEDLTAA FPARAAELIGMHRVPLLS AREVPVPGSLKNVIRVLA | 11.7 | 14.3 |

| | | | | |
|---|---|---|---|---|
| | GNREYLDWLRENTK | LWNTDTPQDRVRHVYLDEAVRLRPDLESPGSHHHHHH | | |
| T33-09.2 | MEEVVLITVPSAEEAVRIAYALVEERLAACVNIIPGLVRIYRWQGRVRVDHVLLLLVKTTTHAFPKLKERVKALHPYTVPEIVALPIAEGNREYLDWLRENTK | MVRGIRGAITVAADTPEAIYAATIELLRRMLEANGIQSYEELAAVIFTVTEDLTAAFPAEAARLIGMHRVPLLSAREVPVPGSLKNVIRVLALWNTDTPQDRVRHVYLGEAVRLRPDLESPGSHHHHHH | 11.8 | 14.3 |
| T33-09.3 | MEEVVLITVPSAEDAVRIARALVEERLAACVNIIPGVVEIYRWQGRVKVKHVLLLLVKTTTHAFPKLKERVKALHPYTVPEIVALPIAEGNREYLDWLRENTK | MVRGIRGAITVKADTPEAIYAATVELLERMLAANGIQSYEELAAVIFTVTEDLTAAFPADAARTIGMHRVPLLSAREVPVPGSLKNVIRVLALWNTDTPQDRVRHVYLGEAVRLRPDLESPGSHHHHHH | 11.7 | 14.2 |
| T33-10.1 | MEEVVLITVPSEEVAIEIAVALVEERLAACVNLVPGLIRIYRWNNRVVIEKELLLLVKTTTHAFPKLKERVKALHPYTVPEIVALPIAEGNREYLDWLRENTKGSHHHHHH | MGAMGPVDEQWIEILRIQALCARYCLTINTQDGEGWAGCFTDDGSFSFDGWTITGRPALREYADAHARVVRGRHLTTDLLYTVIGNIATGRSASVVTLATAAGYKILGSGEYHDLLLKKDGQWRIAHRDLRNDRLVSDPSVAVNVADADVAAVVGHLLAAARRLGTQDKN | 12.8 | 18.4 |
| T33-10.2 | MEEVVLITVPSEGVASEIACALVHERLAACVNIVPGLTRVYRWNNRVKVENELLLLVKTTTHAFPKLKERVKALHPYTVPEIVALPIAEGNREYLDWLRENTKGSHHHHHH | GHMGPVDEQWIEILRIQALCARYCLTINTQDGEGWAGCFTEDGSFSFDGWTITGRPALREYADAHARVVRGRHLTTDLLYTVSGNVASGRSASVVTLATAAGYKILGSGEYHDLLVKKDGQWRIAHRDLRNDRLVSDPSVAVNVADADVAAVVGHLLAAARRLGTQDKE | 12.7 | 18.3 |
| T33-10.3 | MEEVVLITVPSEEVAIEIALALVEERLAACVNIVPGLIRIYRWDNRVVIDKELLLLVKTTTHAFPKLKERVKALHPYTVPEIVALPIAEGNREYLDWLRENTKGSHHHHHH | MGAMGPVDEQWIEILRIQALCARYCLTINTQDGEGWAGCFTEDGSFSFDGWTITGRPALREYADAHARVVRGRHLTTDLLYTVIGNLATGRSASVVTLATAAGYKILGSGEYHDVLLKKDGQWRIASRDLRNDRLVSDPSVAVNVADADVAAVVGHLLAAARRLGTQDES | 12.8 | 18.4 |
| T33-11.1 | MEEVVLITVPSDEEAVTIAATLVSERLAACVNIVPGLTSLYRWNNKVKSEKEYLLLVKTTTHAFPKLKE | MSQAIGILELTSIAKGMELGDAMLKSANVDLLVSKTISPGKFLLMLGGDIGAIQQAIETGVGQAGEMLVDSL | 12.6 | 19.2 |

| | | | | |
|---|---|---|---|---|
| | RVKALHSYTVPEIVALPIAEGNREYLDWLRENTKGSHHHHHH | VLENIHPSVLPAISGLNSVDKRQAVGIVETWSVAACIKAANVALESSDVTLVRVHMAFGIGGKCYMVVAGDVAQVELAVTAASLVAGSRGLLVYRSVIPRPHEAMWRQMVEG | | |
| T33-11.2 | MEEVVLITVPSLEEALVIAGTLVNERLAACVNIVPGLTSLYRWNNEVKSAKEYLLLVKTTTHAFPKLKERVKALHRYTVPEIVALPIAEGNREYLDWLRENTKGSHHHHHH | MSQAIGILELTSIAKGMELGDAMLKSANVDLLVSKTISPGKFLLMLGGDIGAIQQAIETGREQAGEMLVDSLVLEDVHPSVLPAISGLNSVDKRQAVGIVETWSVAACIRAANVALASSDVTLVRVHMAFGIGGKCYMVVAGDVGSVRRAVEAAAIEAGSRGLLVYRSVIPRPHEAMWRQMVEG | 12.7 | 19.3 |
| T33-11.3 | MEEVVLITVPSLHEAYVIAATLVRERLAACVNIVPGLTSLYRWDGVVKVEREYLLLVKTTTHAFPKLKERVKALHSYTVPEIVALPIAEGNREYLDWLRENTKGSHHHHHH | MSQAIGILELTSIAKGMELGDAMLKSANVDLLVSKTISPGKFLLMLGGDIGAIQQAIETGREQAGEMLVDSLVLENIHPSVLPAISGLNSVDKRQAVGIVETWSVAACIRAANVALAGSNVTLVRVHMAFGIGGKCYMVVAGDVAEVEEAVALASEVAGRRGLLVYRSVIPRPHEAMWRQMVEG | 12.7 | 19.3 |
| T33-12.1 | MILVYSTFPNLIEAVRIGIKLLEKRLIACFNAFPITAAYWWKGEIRIDRETAAIFKTTEEKEKELYEELRKLHPYETPAIFTLKVENVLTEYMNWLRESVKGSHHHHHH | MVYMVYVSQDSLTPEAKQAVADAIVDAHRSLTGTQHFLAQVNFQEQPAGNVFLGGVQQGGYTIFVHGLHREGRSADLKGQLAEDIILLVSAAANIDPKHIWVYFGEMPAQQMVEYGRS | 13.0 | 12.9 |
| T33-12.2 | MILVYSTFPDLITAATIGIKLMEKRLIACFNTFPITAVYWWKGEIRVDRETAAIFKTTEEKEKELYEELRKLHPYETPAIFTLKVENVLTEYMNWLRESVKGSHHHHHH | MVYMVYVSQDFLTPEAKAAVARAITDAHRALTGTQHFLAQVNFQEQPAGNVFLGGVQQGGYTIFVHGLHREGRSADLKGQLARDIILLVSVAANIDEKHIWVYFGEMPAQQMVEYGRE | 12.9 | 13.1 |
| T33-13.1 | MHNFIYITASSAEEAVEIAVRLLEKKLAACVNLYPIVELFWWEGRIRAEQEVAMIVKTRSEKFAEVRDEVKAMHSYTTPCICAIPIERGLKEFLDWIDETVE | MVRGIRGAITVKRNTPLAIYASTVALLREMLEANGIQSYEELAAVIFTVTEDLTAASPADAARAIGMHRVPLLSAREVPVPGSLKNVIRVLALWNTDTPQDRVRHVYLGEAVRLRPDLESPGSHHHHHH | 11.8 | 14.2 |
| T33-13.2 | MHNFIYITAKSGVEAVGIAYRLLEKKLAACVNLYPI | MVRGIRGAITVGRDTPLAISAATIALLKEMLEANGIQ | 11.7 | 14.1 |

| | | | | |
|---|---|---|---|---|
| | VELFWWEGRVHAETEVAMIVKTRSEKFAEVRDEVKAMHSYTTPCICAIPIERGLKEFLDWIDETVN | SYEELAAVIFTVTEDLTAADPADAARAIGMHRVPLLSAREVPVPGSLKNVIRVLALWNTDTPQDRVRHVYLSEAVRLRPDLESPGSHHHHHH | | |
| T33-13.3 | MHNFIYITADSGIEAVAIAMRLLEKKLAACVNIYPIVELFWWEGEIRVRQETAMIVKTRSEKFAEVRDEVKAMHSYTTPCICAIPIERGLKEFLDWIDETVY | MVRGIRGAITVARDTEIEIAAATMVLLKRMLKANGIQSYEELAAVIFTVTEDLTAANPADAARAIGMHRVPLLSAREVPVPGSLKNVIRVLALWNTDTPQDRVRHVYLDEAVRLRPDLESPGSHHHHHH | 11.9 | 14.2 |
| T33-14.1 | MKIRIGHGFDVHKFGEPRPLILCGVEVPYETGLVAHSDGDVVLHAISDAILGAMALGDIGKHFPDTDAAYKGADSIDLLRECVELARAKGFELGNLDVTIIAQAPKMAPYIKLMQLNLSIVLDADIADINVKATTTEKLGFTGRKEGIAVEAVVLLSRK | MSEKKAVIGVVTISDRASKGIYKDYSGEAIILYLKTVIITPFEVEYRVIPDERDLIEKTLIELADEKGCSLILTTGGTGPAPRDVTPEATEAVCEKMLPGFGELMRQVSLKQVPTAILSRQTAGIRGSCLIVNLPGDPESILLCLKAVMPAIPYCIDLIGGAYIDTDPRIVKAFRPKPGSHHHHHH | 17.0 | 20.3 |
| T33-14.2 | MKIRIGHGFDVHKFGEPRPLILCGVEVPYETGLVAHSDGDVVLHAISDAILGAMALGDIGKHFPDTDAAYKGADSLDLLRHCIDLARAKGFELGNLDVTIIAQAPKMAPYIKLMQLNLSVLLDADIADINVKATTTEKLGFTGRKEGIAVEAVVLLSRK | MSEKKAVIGVVTISDRASKGIYKDYSGLTIINLLKTVIITPFEVEYRVIPDERDLIEKTLIELADEKGCSLILTTGGTGPAPRDVTPEATEAVCEKMLPGFGELMRQVSLKQVPTAILSRQTAGIRGSCLIVNLPGDPESIVICLKAVMPAIPYCIDLIGGAYIDTDPRIVKAFRPKPGSHHHHHH* | 17.0 | 20.2 |
| T33-14.3 | MKIRIGHGFDVHKFGEPRPLILCGVEVPYETGLVAHSDGDVVLHAISDAILGAMALGDIGKHFPDTDAAYKGADSLDLLRECIDLARAKGFELGNLDVTIIAQAPKMAPYIKLMQLNLSVLLDADAADINVKATTTEKLGFTGRKEGIAVEAVVLLSRK | MSEKKAVIGVVTISDRASKGIYKDYSGEAIILLLKTTIITPFEVEYRVIPDERDLIEKTLIELADEKGCSLILTTGGTGPAPRDVTPEATEAVCEKMLPGFGELMRQVSLKQVPTAILSRQTAGIRGSCLIVNLPGDIDSIVLCLKAVMPAIPYCIDLIGGAYIDTDPRIVKAFRPKPGSHHHHHH | 17.0 | 20.2 |
| T33-15.1 | MVRGIRGAVSVLEDTRLVISHATRLLLERMLKANGIQSYEELAAVIFTVTEDLTSAFPAEAARRIGMHRVPLLSAREVPVPGSLPRVIRVLALWNTDTPQDRVRHVYLDDAVRLRPDLES | MSKAKIGIVTVSDRASAGIEEDVSGQAIIDWLKAYLTSEWEPIYQVIPDEQDVIETTLIKMADEQDCCLIVTTGGTGPAKRDVTPEATEAVCDRMMPGFGELMRATLLRFNPTAILSRQTAGLRG | 14.5 | 19 |

| | | | | |
|---|---|---|---|---|
| | PGSHHHHHH | DSLIVNLPGAPDSIILCLE AVFPAIPYCIDLMEGPYL ECDESVLKPFRP | | |
| T33-16.1 | MSLILVYSTFPNRTEAVT IGVKLLKKRLIACFNAFPI VSAYEEDGVIELKEEWA AIFKTTEEKEKELYEELR KLHPYETPAIFTLKVENV LTEYMNWLRESV | MPVIQTFVSTPLDKEKRE ALALEYRVITETVLGKDP ALVMMTFHDSTPMFFNG NTDPVACVRVEALGGYG PSEPEKVTRLVTKAITDIC GIVADRIFVLYFSPLHCG WNGTNLGSHHHHHH | 12.0 | 13.4 |
| T33-16.2 | MSLILVYSTFPNLHEAVV IGVELLRKRLIACFNAFPI VSVYEEDGVIVVREEWA AIFKTTEEKEKELYEELR KLHPYETPAIFTLKVENV LTEYMNWLRESI | MPVIQTFVSTPLDEEKRE ALALQYRVVTYTILKKEP DLVMMTFHDSTPMRFRG STAPVACVRVEALGGYG PSEPERVTRVVTAAITAV CGIVADRIFVLYFSPLHC GWNGTNLGSHHHHHH | 12.0 | 13.5 |
| T33-17.1 | MSLILVYSTFPDLVSAIAI GIKLLEKRLIACFNAFPIT SVYWWKGEIRVEKETA AIFKTTEEKEKELYEELR KLHPYETPAIFTLKVENV LTEYMNWLRESVGSHH HHHH | MVYMVYVSQGRLTPEQK AAVARAIVDAHRALTGTQ HFLAQVNFQEQPKGNVF LGGKRQDGNTIFVHGLH REGRSADLKGQLARDIIL LVSAAADIPEKHIWVYFG EMPAQQMVEYGRE | 13.0 | 13.2 |
| T33-17.2 | MSLILVYSTFPSLTEAVA IGIELIEKRLIACFNAFPIT SVYWWKGEIRVERETA AIFKTTEEKEKELYEELR KLHPYETPAIFTLKVENV LTEYMNWLRESVGSHH HHHH | MVYMVYVSQGQLSPEQ KAAVADAIVQAHRRLTGT QHFLAQVNFQEQPKGNV FLGGVRQDGNTIFVHGL HREGRSADLKGQLAHDII LLVSAAANIPEKHIWVYF GEMPAQQMVEYGRA | 13.0 | 13.1 |
| T33-17.3 | MSLILVYSTFPNRVTAIAI GLKLIEKRLIACFNAFPIT TVYWWKGEIRVERETA AIFKTTEEKEKELYEELR KLHPYETPAIFTLKVENV LTEYMNWLRESVGSHH HHHH | MVYMVYVSQGYLTPQQK ADVAAAIVTAHRDLTGTQ HFLAQVNFQEQPKGNVF LGGVRQDGNTIFVHGLH REGRSADLKGQLARDIIL LVSAAANIPEKHIWVYFG EMPAQQMVEYGRS | 13.1 | 13.1 |
| T33-18.1 | MSLILVYSTFPNRLEALLI GLELLEKRLIACFNAFEI TSAYWEKGRIRIEREWA AIFKTTEEKEKELYEELR KLHPYETPAIFTLKVENV LTEYMNWLRESVGSHH HHHH | MVYMVYVSQDRLTPSAK HAVAQAITDAHLAHTGEE HTLAQVNFQEQPAGNVF LGGVQQGGDTIFVHGLH REGRSDELKRDLITDIQV LVSHAANIDPKHIWVYFG EMPASQMVEYGGL | 13.2 | 13 |
| T33-18.2 | MSLILVYSTFPNLLEAKLI GLKLLKKRLIACFNAFEI TSAYWEKGRIRTRREW AAIFKTTEEKEKELYEEL RKLHPYETPAIFTLKVEN VLTEYMNWLRESVGSH HHHHH | MVYMVYVSQDRLTPSAK HAVAQAITDAHLTHTGEE HSLAQVNFQEQPAGNVF LGGVQQGGDTIFVHGLH REGRSDELKQRLITDIIAK VSIAADIDPKHIWVYFGE MPASQMVEYGGL | 13.2 | 13 |

| | | | | |
|---|---|---|---|---|
| T33-18.3 | MSLILVYSTFPNLLEALYIGLLLLDKRLIACFNAFEIVSAYWEKGRIRIRREWAAIFKTTEEKEKELYEELRKLHPYETPAIFTLKVENVLTEYMNWLRESVGSHHHHHH | MVYMVYVSQDRLTPSAKHDVARAITDAHLAFTGEEHRLAQVNFQEQPAGNVFLGGVQQGGDTIFVHGLHREGRSDELKRDLITAIIAHVSLAANIDPKHIWVYFGEMPRVQMMEYGGL | 13.2 | 13.2 |
| T33-19.1 | MDSPIIEANGTLDELTSFIGEAKHYVDEEMKGILEEIQNDIYKIMGEIGSRGKIEGISPERLRWLLDLIDRYSEMVKKENVLPGGTLESAKLDVCRTIAERAARKVKTVVEEKGIGETALHYLEVLSLLLELLARVIEIEKNKEGSHHHHHH | MKKIIETQRAPGAIGPYVQGVDLGSMVFTSGQIPVPKTGKIPSTIAEQARLSLENVKAIVVAAGLSVGDIIKMTVFITDLSDLPLIELVYERFFDEHQATYPTRSCVQVARLPKDVKLEIEAIAVRD | 17.2 | 13.8 |
| T33-19.2 | MDSPIIEANGTLDELTSFIGEAKHYVDEEMKGILEEIQNDIYKIMGEIGSRGKIEGISEERLLFLLDLIDRYSEMVPLSGVLPGGTLESAKLDVCRTIAERAARKVKTVVEETGIGEVALRYLEVLSLLLELLARVIEIEKNKEGSHHHHHH | MKKIIETQRAPGAIGPYVQGVDLGSMVFTSGQIPVPETGEIPEHVYEQARLSLENVKAIVVAAGLSVGDIIKMTVFITDSSDLPIIELVYKRFFDEHQATYPTRSCVQVARLPKDVKLEIEAIAVRP | 17.0 | 14.0 |
| T33-19.3 | MDSPIIEANGTLDELTSFIGEAKHYVDEEMKGILEEIQNDIYKIMGEIGSRGKIEGISEERLDELLDLYNRYSEMVPRDNVLPGGTLESAKLDVCRTIAERAARKVKTVVEETGIGRVALQYLEVLSLLLELLARVIEIEKNKEGSHHHHHH | MKKIIETQRAPGAIGPYVQGVDLGSMVFTSGQIPVPETGRIPAHVAEQARLSLENVKAIVVAAGLSVGDIIKMTVFITDLEDLPLIEAVYKRFFDSHQATYPTRSCVQVARLPKDVKLEIEAIAVRE | 17.2 | 13.8 |
| T33-20.1 | MSPIEEANGTLDELTSFIGEAKHYVDEEMKGILEEIQNDIYKIMGEIGSKGKIPGIPQSRLDWLLDLIERYKEMVNRKFVLPGGTLESAKLDVCRTIARRATRKVLKVVEEFGIGRVAVQYLLVLVELLFLLARVIEIEKNKEGSHHHHHH | MLYFQGMPHLVIEATANLRLETSPGELLEQANAALFDSGQFGEADIKSRFVTLEAYRQGTAAVERAYLHACLSILDGRSVLERTRLALRLRAVLAGAVAGGGEEGVQVSVEVREMERISYAKAVVAR | 17.2 | 13.8 |
| T33-20.2 | MKPIEEANGTLDELTSFIGEAKHYVDEEMKGILEEIQNDIYKIMGEIGSKGKIPGIPESRLEWLLSLIARYSEMVNREFVLPGGTLESAKLDVCRTIARRAARKVQEVLEEFGIGDVALKYLLVLVELLFLLARVIEIEKNKEGSHHHHHH | MLYFQGMPHLVIEATANLRLETSPGELLEQANAALFASGQFGEADIKSRFVTLEAYRQGTAAVERAYLHACLSILDGRSAETRLRLSIRLCAVLASAVAGGGEEGVQVSVEVREMERISYAKAVVAP | 17.1 | 13.7 |
| T33-20.3 | MEPIEEANGTLDELTSFI | MLYFQGMPHLVIEATANL | 17.2 | 13.7 |

| | | | | |
|---|---|---|---|---|
| | GEAKHYVDEEMKGILEEIQNDIYKIMGEIGSKGKIPGISEDRLDWLLDLIDRYKEMVNRKFVLPGGTLESAKLDVCRTIAKRAARKVLKVLEEFGIGDVALKYLLVLEELLFLLARVIEIEKNKEGSHHHHHH | RLETSPGELLEQANAALFDSGQFGEADIKSRFVTLEAYRQGTAAVERAYLHACLSILDGRSTETRLRLSLRLLAVLAGAVAGGGEEGVQVSVEVREMERISYAKAVVAA | | |
| T33-21.1 | MDSPIIEANGTLDELTSFIGEAKHYVDEEMKGILEEIQNDIYKIMGEIGSKGKIPGIPQSSLARLWDLIKRYSEMVNKSFVLPGGTLESAKLDVCRTIARRAARKVATVVEEFGIGGVALAYLDVLSELLFLLARVIEIEKNKEGSHHHHHH | MLDFQGMPHLVIEATANLRLETSPGELLEQANAALFASGQFGEADIKSRFVTLEAYRQGTAAVERAYLHACLSILDGRSIGELRALGARLCAVLAAAVAGGGEEGVQVSVEVREMERLSYAKRVVAA | 16.9 | 13.5 |
| T33-21.2 | MDSPIIEANGTLDELTSFIGEAKHYVDEEMKGILEEIQNDIYKIMGEIGSKGKIPGISEESLVKLFDLIKRYSEMVNKSFVLPGGTLESAKLDVCRTIARRAARKVATVVEEFGIGAVALAYLELLSELLFLLARVIEIEKNKEGSHHHHHH | MLDFQGMPHLVIEATANLRLETSPGELLEQANRALFDSGQFGEADIKSRFVTLEAYRQGTAAVERAYLHACLSILDGRDIGTRTALAARLLAVLAGAVAGGGEEGVQVSVEVREMERLSYAKRVVAA | 16.9 | 13.6 |
| T33-21.3 | MDSPIIEANGTLDELTSFIGEAKHYVDEEMKGILEEIQNDIYKIMGEIGSKGKIPGISYESIARLFDLIRRYKEMVNKSFVLPGGTLESAKLDVCRTIARRAARKVATVLEEFGIGAVALAYLELLSELLFLLARVIEIEKNKEGSHHHHHH | MLDFQGMPHLVIEATANLRLETSPGELLEQANAALFDSGQFGEADIKSRFVTLEAYRQGTAAVERAYLHACLSILDGRSTGELTALAARLCAVLAGAVAGGGEEGVQVSVEVREMERLSYAKRVVAA | 17.1 | 13.5 |
| T33-22.1 | MDSPIIEANGTLDELTSFIGEAKHYVDEEMKGILEEIQNDIYKIMGEIGSKGKIEGISEDRLADLLELLLRYSAMVNKSFVLPGGTLESAKLDVCRTIARRAERKVATVLREFGIGKVALRYLRVLERLLFLLARVIEIEKNKEGSHHHHHH | MSQAIGILELTSIAKGMELGDAMLKSANVDLLVSKTISPGKFLLMLGGDYGAIQQAIETGTSQAGEMLVDSELLKNIHPSVLPAISGLNSVDKRQAVGIVETWGVTACIVAADFAVKNSNVTLVRVHMAFGIGGKCYMVVAGDVSDVNNAVDVASAVAGALGKLVYRSVIPRPHEAMWRQMVEG | 17.2 | 19.2 |
| T33-22.2 | MDSPIIEANGTLDELTSFIGEAKHYVDEEMKGILEEIQNDIYKIMGEIGSKGKIEGISEERLVDLLELLLRYSKMVNKSFVLPGGTLESAKLDVCRTIARRAERKVATVLREFGIGKVALRYLRVLERLLFLLARVIEIEKN | MSQAIGILELTSIAKGMELGDAMLKSANVDLLVSKTISPGKFLLMLGGDYGAIQQAIETGTSQAGEMLVDSALLKDIHPSVLPAISGLNSVDKRQAVGIVETWGVEACIIAADFAVKGSNVTLVRVHMAFGIGGKCYMVVAG | 17.3 | 19.2 |

| | | | | |
|---|---|---|---|---|
| | KEGSHHHHHH | DVSDVNNAVEVASAVAGALGRLVYRSVIPRPHEAMWRQMVEG | | |
| T33-22.3 | MDSPIIEANGTLDELTSFIGEAKHYVDEEMKGILEEIQNDIYKIMGEIGSKGKIEGISEDRVAYLLELLLRYEKMVNKSFVLPGGTLESAKLDVCRTIARRAERKVATVLREFGIGKVALRYLKVLERLLFLLARVIEIEKNKEGSHHHHHH | MSQAIGILELTSIAKGMELGDAMLKSANVDLLVSKTISPGKFLLMLGGDTGAIQQAIETGTSQAGEMLVDSDLIKDIHPSVLPAISGLNSVDKRQAVGIVETWGVTACIIAADFAVKGSNVTLVRVHMAFGIGGKCYMVVAGDVSDVNNAVDVASRVAGALGLLVYRSVIPRPHEAMWRQMVEG | 17.3 | 19.2 |
| T33-23.1 | MDSPIIEANGTLDELTSFIGEAKHYVDEEMKGILEEIQNDIYKIMGEIGSKGKIPGISISSYVWLVGLIRRYEEMVNKSFVLPGGTLESAKLDVCRTIARRAHRKVKTVVEEFGIGEDAAAYLELLSRLLFLLARVIEIEKNKEGSHHHHHH | MSQAIGILELTSIAKGMELGDAMLKSANVDLLVSKTISPGKFLLMLGGDIGAIQQAIETGTSQAGEMLVDSLVIPDIHPSVLPAISGLNSVDKRQAVGIVETWSVVACIRAANAAVASSNVTLVRVHMAFGIGGKCYMVVAGDVSDVNEAVTVASAVAGATGLLVYRSVIPRPHEAMWRQMVEG | 17.1 | 19 |
| T33-23.2 | MDSPIIEANGTLDELTSFIGEAKHYVDEEMKGILEEIQNDIYKIMGEIGSKGKIPGIPWESLLWLRDLIKRYEEMVNKSFVLPGGTLESAKLDVCRTIARRAYRKVKTVVEEFGIGEVAVQYLRELSRLLFLLARVIEIEKNKEGSHHHHHH | MSQAIGILELTSIAKGMELGDAMLKSANVDLLVSKTISPGKFLLMLGGDIGAIQQAIETGTSQAGEMLVDSLVLPDIHPSVLPAISGLNSVDKRQAVGIVETWSVEACIRAADMAVRTSNVTLVRVHMAFGIGGKCYMVVAGDVSDVNEAVTAASATAGALGKLVYRSVIPRPHEAMWRQMVEG | 17.5 | 19.2 |
| T33-23.3 | MDSPIIEANGTLDELTSFIGEAKHYVDEEMKGILEEIQNDIYKIMGEIGSKGKIPGIPLSSYIWLLELIRRYEEMVNKSFVLPGGTLESAKLDVCRTIARRAFRKVRTVVEEFGIGEVAAVYLRVLSELLFLLARVIEIEKNKEGSHHHHHH | MSQAIGILELTSIAKGMELGDAMLKSANVDLLVSKTISPGKFLLMLGGDIGAIQQAIETGTSQAGEMLVDSLVIPDIHPSVLPAISGLNSVDKRQAVGIVETWSVTACIIAANAAVAGSNVTLVRVHMAFGIGGKCYMVVAGDVSDVNEAVTVASLTAGALGRLVYRSVIPRPHEAMWRQMVEG | 17.3 | 19.1 |
| T33-24.1 | MPLLKFDLFYGRSDEQIKSLIDAAHAAMVLAFGVAPTDRYQTVSQHREGEMVLEDTGLGYGRTEAVVLLTVISRPRSEEQKVLFNRLLVAALESVCGISPDDVIVALVENSDADWSFG | MSKAKIGIVTVSDRAYAGIYEDISGKAIIDTLNDYLTSEWEPIYRVIPDNLEQIKVALGYMALIEDCCLIVTTGGTGPAKRDVTPEATEAVCDRMMPGFGELMRAESLKFVPTAILSRQTAGLLGDS | 15.2 | 19 |

| | | | | |
|---|---|---|---|---|
| | GGRAEFLTGDLVGGSH HHHHH | LIVNLPGKPKSIRECLDAV FPAIPYCIDLMEGPYLEC NEAVIKPFRP | | |
| T33-24.2 | MPLLKFDLFYGRSDEQI KSLIDAAWAAMVLAFGV PATDRYQTVSQHREGE MVLEDTGLGYGRTRAV VLLTVISRPRSEEQKVLF NRLLCAALEVVCGISPD DVIVALVENSDADWSFG RGRAEFLTGDLVGGSH HHHHH | MSKAKIGIVTVSDRAFAGI YEDISGKAIIDTLNDYLTS EWEPIYRVIPDELGLIEAA LAYMALVEDCCLIVTTGG TGPAKRDVTPEATERVC DRMMPGFGELMRAESLK FVPTAILSRQTAGLLGDS LIVNLPGKPKSIRECLDAV FPAIPYCIDLMEGPYLEC NEAVIKPFRP | 15.4 | 19.0 |
| T33-24.3 | MPLLKFDLFYGRSDEQI KSLIDAAHGAMVLAFGV PASDRYQTVSQHRPGE MVLEDTGLGYGRTDAV VLLTVISRPRSEEQKVLF NRLLTAALEVLCGISPDD VIVALVENSDADWSFGG GRAEFLTGDLVGGSHH HHHH | MSKAKIGIVTVSDRAFAGI YEDISGKAIIDTLNDYLTS EWEPIYRVVPDDKDIIVTT LAYMALIEDCCLIVTTGGT GPAKRDVTPEATEAVCD RMMPGFGELMRAESLKF VPTAILSRQTAGLLGDSLI VNLPGKPKSIRECLDAVF PAIPYCIDLMEGPYLECN EAVIKPFRP | 15.2 | 19.0 |
| T33-25.1 | MSRVYLIFSTCPDLPSA EIISRVLVQERLAACVTQ LPGAVSTYRWQGKIETT QEIQLLIKTNRKTVALAM LKLKDLHPYRLPETIAVQ VSTGYPRFEKWIEDEIE | MSRTMVSSGSLFEEIFGY SRAVRIGPLVVVAGTTGS GSTIGAQTLDALRRIEIAL GQAGATLADVVRTRIYVT DISLYDVVGMAIYDAFRKI RPVTSMVEVTALIAPGLL VEIEADAYGGSHHHHHH | 12.0 | 13.6 |
| T33-25.2 | MSSVYLIFSTCPDLPSA EIISRVLVQERLAACVTQ LPGAVSTYRWQGKIETT QEIQLLIKTNRRTVSLAIL KLQDLHPYRLPETIAVQ VSTAYPEFEKWIYDEIE | MSRTMVSSGSLLEEVTG YSRAVRIGPLVVVAGTTG SGKDIGAQTLDALRRIEIA LGQAGATLADVVRTRIYV TDISRFDDVALAINRAFR KIRPVTSMVEVTALIAPGL LVEIEADAYGGSHHHHH H | 12.0 | 13.5 |
| T33-25.3 | MSNVYLIFSTCPDLPSA EIISRVLVQERLAACVTQ LPGAVSTYRWQGKIETT QEIQLLIKTNRDTIVRAIL KLKELHPYRLPETIAVQV ATAYAGFEKWIWDEIA | MSRTMVSSGSLLEERLG YSRAVRIGPLVVVAGTTG SGDTIGMQTLDALRRIEIA LGQAGATLADVVRTRIYV TDISRIDDVGLAIAEAFGK IRPVTSMVEVTALIAPGLL VEIEADAYGGSHHHHHH | 11.9 | 13.4 |
| T33-26.1 | MKSELEKMLAGHLYNP ADPELQDLVLLARRLVD HYNRTSADEYKERQTLL RALFGSTGERLFIEPNF RCDYGENIHVGENFFM NFDGVILDVCEVRIGDH CFIGPGVHIYTATHPLDP HERNSGLEYGKPVVIGH | MPVIQTFVSTPLDERTRS LLAAVYARVTREVLGKDP TRVMMTFHDSTPMHHK GSTAPVACVRVEALGGY GPSEPEKVTSIVTRAITDL CGIVADRIFVLYFSPLHC GWNGTNV | 21.3 | 12.4 |

| | | | | |
|---|---|---|---|---|
| | NVWIGGRAVINPGVTIG DNAVIASGAVVTKDVPA NAVVGGNPAKVIKWLG GSHHHHHH | | | |
| T33-26.2 | MKSELEKMLAGHLYNP ADPELQDLRLLARRLVD AYNETSADEYEERKLLL DTLFGSTGERLFIEPNF RCDYGDNIHVGENFFM NFDGVILDVCEVRIGDH CFIGPGVHIYTATHPLDP HERNSGLEYGKPVVIGH NVWIGGRAVINPGVTIG DNAVIASGAVVTKDVPA NAVVGGNPAKVIKWLK GSHHHHHH | MPVIQTFVSTPLDERDRL LLAAVYARVTEKVLGKDP SKVMMTFHDSTPMHHR GSTAPVACVRVEALGGY GPSEPEKVTSIVTDAITEV CGIVADRIFVLYFSPLHC GWNGTNV | 21.3 | 12.3 |
| T33-26.3 | MKSEKEKMLAGHLYNP ADKELQNELLTARRLVD LYNETGADEYDERRVLL RTLFGSTGERLFIEPNF RCDYGRNIHVGENFFM NFDGVILDVCEVRIGDH CFIGPGVHIYTATHPLDP HERNSGLEYGKPVVIGH NVWIGGRAVINPGVTIG DNAVIASGAVVTKDVPA NAVVGGNPAKVIKWLG GSHHHHHH | MPVIQTFVSTPLDERERT LLAAVYARVTREVLGKPS EKVMMTFHDSTPMHHN GSTAPVACVRVEALGGY GPSEPEKVTSIVTAAITDV CGIVADRIFVLYFSPLHC GWNGTNV | 21.3 | 12.3 |
| T33-27.1 | MTMADETIILNVLGQYT RAHDRRDPDAMAALFA PDASIVVLDAVGGASKPI SVLHGRDAIRVAVRQM MAPHGYRAWSQNVVN APVIHIHGDTARLDAQF MVFSILAAEVPDGGWPT GTFGAQGRIVPIEAGTY TLFLRTVPDGWVIAHMV IKHRLPMAFG | MSQAIGILELSSIAKGMEL GDAMLKAANVDLLVSKTI SPGKFLLMLGGDLDDIILA VAVGMERAGDSLLDSEVI PDIHPSVLPAISGLNSVD KRQAVGIVETWSVAACIK AADRAVKGSNVTLVRVH MAFGIGGKCYMVVAGDV SDVNNAVTVASESAGEK GLLVYRSVIPRPHEAMW RQMVEGGSHHHHHH | 17.3 | 20.2 |
| T33-27.2 | MTMADETIILNVLGQYT RAHDRRDPDAMAALFA PDATIVVVDAVGGANRII SLLDGRDAIRVAVRQM MAPHGYRAWSQNVVN APVIHINGDKALLDAQF MVFSILAAEVPDGGWPT GTFGAQGRIVPIEAGEY LLMLETVPDGWVISSMII KHRLPMAFG | MSQAIGILVLSSIAKGMEL GDAMLKAANVDLLVSKTI SPGKFLLMLGGDESAIKQ AVAVGVERAGDALLDSA VISDIHPSVLPAISGLNSV DKRQAVGIVETWSVAACI EAADRAVKGSNVTLVRV HMAFGIGGKCYMVVAGD VSDVNNAVTVASESAGE KGLLVYRSVIPRPHEAM WRQMVEGGSHHHHHH | 17.3 | 20.0 |
| T33_27.3 | MTMADETIILNVLGQYT RAHDRRDPDAMAALFA PDATIVVVDAVGGAFRVI SILKGRDAIRVAVRQMM | MSQAIGILVLSSIAKGMEL GDAMLKAANVDLLVSKTI SPGKFLLMLGGDEGAIK QAVAVGVRNAGDDLLDS | 17.4 | 20.1 |

| | | | | |
|---|---|---|---|---|
| | APHGYRAWSQNVVNAPVIHIKGDKALLDAQFMVFSILAAEVPDGGWPTGTFGAQGRIVPIEAGTYLLMLETVEDGWVISRMIIEHRLPMAFG | KVIDNIHPSVLPAISGLNSVDKRQAVGIVETWSVAACIRAADRAVKGSNVTLVRVHMAFGIGGKCYMVVAGDVSDVNNAVTVASESAGEKGLLVYRSVIPRPHEAMWRQMVEGGSHHHHHH | | |
| T33-28.1 | MSVNTSFLSPSLVTIRDFDHGQFAVLRIGRTGFPADKGDIDLCLSKMQGVLSAQLFLGNPREPGFKGPHIRIRCVDIDDKHTYNAMVYVDLIVGTGASEVERETAEEKARAALAVALRVDEADEHSCVTQFEMKLREELLSSDSFHPDKDEYYKDFL | MPVIQTFVSTPLDHEKRTLLFRQYRIVTAVILGKPAELVMMTFHDSTPMHFFGSTDPVACVRVEALGGYGPSEPEKVTEVVTKAISYVCGIVADRIFVLYFSPLHCGWNGTNVGSHHHHHH | 17.6 | 13.5 |
| T33-28.2 | MSVNTSFLSPSLVTIRDFDNGQFAVLRIGRTGFPADKGDIDLCLRKMEGVLAAQIYLGNPREPGFKGPHIRIRCVDIDDKHTYNAMVYVDLIVGTGASEVERETAEELAKAALDIALEVDKANEHSCVTQFEMKLREELLSSDSFHPDKDEYYKDFL | MPVIQTFVSTPLDHRKREMLSTVYRIVTATILGKPPELVMMTFHDSTPMHFFGSTDPVACVRVEALGGYGPSEPEKVTKVVTEAISYLCGIVADRIFVLYFSPLHCGWNGTNIGSHHHHHH | 17.6 | 13.5 |
| T33-28.3 | MSVNTSFLSPSLVTIRDFDKGQFAVLRIGRTGFPADKGDIDLCLSKMDGVLAAQLYLGNPREPGFKGPHIRIRCVDIDDKHTYNAMVYVDLIVGTGASEVERETAEERARRALAVALRVDEADEHSCVTQFEMKLREELLSSDSFHPDKDEYYKDFL | MPVIQTFVSTPLDHEKRNMLTKVYRIVTDTILGKPAELVMMTFHDSTPMHFFGSTDPVACVRVEALGGYGPSEPEKVTKVVTDAISYVCGIVADRIFVLYFSPLHCGWNGTNLGSHHHHHH | 17.7 | 13.5 |
| T33-29.1 | MSRTMVSSGSRYERIMGYSRAVRIGPLVVVAGTTGSGRGIGDQTEDALRRIEIALGQAGATLADVVRTRIYVTDISEFAAVAIQHYVAFRKIRPVTSMVEVTALIAPGLLVEIEADAYGGSHHHHHH | MKKIEAIIRPFKLDEVKIALVNAGIVGMTVSEVRGFGRQKRGSEYTVEFLQKLKLEIVVLDEDVAEVILKIREAARTGENGDGKIFVSPVLRVVRIRDGAMDEAAISAWA | 13.7 | 12.2 |
| T33-29.2 | MSRTMVSSGSKEEEIFGYSRAVRIGPLVVVAGTTGSGRTIAAQTEDALRRIEIALGQAGATLADVVRTRIYVTDISRWDEVGLVHKNAFAKIRPVTSMVEVTALIAPGLLVEIEADAYGGSHHHHHH | MKKIEAIIRPFKLDEVKIALVNAGIVGMTVSEVRGFGRQKRGSEYTVEFLQKLKLEIVVLDEDVPLVINKIREAARTGENGDGKIFVSPVERVVRIRDGAMDELAISAWS | 13.7 | 12.3 |

| | | | | |
|---|---|---|---|---|
| T33-30.1 | MSKAKIGIVTVSDRASA GTLLDTNGLAIRSCLDM YLTSEWEPIYQVIPDEQ DVIETTLIKMADEQDCCL IVTTGGTGPAKRDVTPE ATEAVCDRMMPGFGEL MRAESLKFVPTAILSRQ TAGLRGDSLIVNLPGSP ESIFECLKAVFPAIPYCID LMEGPYLECNERVIKPF RPGSHHHHHH | MPFLELDTNLPANRVPA GLEKKLCEQAAAILGKPA DRVNVTVRPGLAMALSG STEPCAQLSISSIGVVGE AERNNLISRGFTDFLTKE LALGQDRILIRFFPLESW QIGKIGLVMTFE | 20.0 | 12.8 |
| T33-30.2 | MSKAKIGIVTVSDRASA GTLLDTNGLAIRTALRRY LTSEWEPIYQVIPDEQD VIETTLIKMADEQDCCLI VTTGGTGPAKRDVTPE ATEAVCDRMMPGFGEL MRAESLKFVPTAILSRQ TAGLRGDSLIVNLPGSP ESIIECLKAVFPAIPYCID LMEGPYLECNEKVIKPF RPGSHHHHHH | MPFLELDTNLPANRVPA GLEKRLCEVAAEILGKPA DRVNVTVRPGLAMALSG STEPCAQLSISSIGVVGT AERNAVISAGFTDFLTKE LALGQDRILIRFFPLESW QIGKIGLVMTFD | 20.0 | 12.7 |
| T33-30.3 | MSKAKIGIVTVSDRASA GVRLNRNGLAIETWLDL YLTSEWEPIYQVIPDEQ DVIETTLIKMADEQDCCL IVTTGGTGPAKRDVTPE ATEAVCDRMMPGFGEL MRAESLKFVPTAILSRQ TAGLRGDSLIVNLPGDP DSIIECLKAVFPAIPYCID LMEGPYLECNEKVIKPF RPGSHHHHHH | MPFLELDTNLPANRVPA GLEKKLCRAAAEILGKPE DRVNVTVRPGLAMALSG STEPCAQLSISSIGVVGT AERNALISRRFTDFLTKE LALGQDRILIRFFPLESW QIGKIGLVMTFD | 20.1 | 12.9 |

**Table S3. Amino acid sequence of individual T33 components for *in vitro* assembly.**

| Design | Sequences | MW |
|---|---|---|
| T33-06.1A | MHQIRVGVLTVSDSCFRNL<br>RPDLSGRALERYVQDPKLL<br>GGTISAYKIVPDEIEEIKETL<br>IDWCDEKELNLILTTGGTG<br>FAPRDVTPEATKEVIEREA<br>PGMALAMLMGSLNITPLG<br>MLSRPVCGIRGKTLIINLPG<br>SLLGSLRCFDFILPALPHAI<br>DLLRDAIVKVKEVHHGSHH<br>HHHH | 19.6 |
| T33-06.1B | MPVIQTFVSTPLDEDDRRA<br>LSLVYRYATEKILGKPADLV<br>MMTFHDSTPMHFFGSTDP<br>VACVRVEALGGYGPSEPE<br>EVTKLVTAAITEVCGIVADR<br>IFVLYFSPLHCGWNGTNVG<br>SHHHHHH | 13.4 |
| T33-08.2A | MSPVVEVQGTIDELNSFIG<br>YALVLSRWDDIRNDLFRIQ<br>NDLFVLGEDVSTGGKGRT<br>VTLEMIAELVKKSYKMKKEI<br>GKIELFVVPGGSVESASLH<br>MARAVSRRLERRIEAAAKL<br>TEINELVLLYAQALSRILFM<br>HALISNKRLNIPEKIYGSHH<br>HHHH | 17.9 |
| T33-08.2B | MNQPIIEANGTLDELTSFIG<br>EAKHYVDEEMKGILEEIQN<br>DIYKIMGEIGSKGKIEGISD<br>ESLVKLLDLIERYEEMVNK<br>SFVLPGGTLESAKLDVCRT<br>IARRATLKVKTVLEEFGIGF<br>NAVLYLEVLSELLFLLARVI<br>EIEKNKEGSHHHHHH | 17.2 |
| T33-11.1A | MEEVVLITVPSDEEAVTIAA<br>TLVSERLAACVNIVPGLTSL<br>YRWNNKVKSEKEYLLLVKT<br>TTHAFPKLKERVKALHSYT<br>VPEIVALPIAEGNREYLDW<br>LRENTKGSHHHHHH | 12.6 |
| T33-11.1B | MSQAIGILELTSIAKGMELG<br>DAMLKSANVDLLVSKTISP<br>GKFLLMLGGDIGAIQQAIET<br>GVGQAGEMLVDSLVLENIH<br>PSVLPAISGLNSVDKRQAV | 20.1 |

| | | |
|---|---|---|
| | GIVETWSVAACIKAANVAL ESSDVTLVRVHMAFGIGGK CYMVVAGDVAQVELAVTA ASLVAGSRGLLVYRSVIPR PHEAMWRQMVEGGSHHH HHH | |
| T33-18.2A | MSLILVYSTFPNLLEAKLIG LKLLKKRLIACFNAFEITSA YWEKGRIRTRREWAAIFKT TEEKEKELYEELRKLHPYE TPAIFTLKVENVLTEYMNW LRESVGSHHHHHH | 13.2 |
| T33-18.2B | MVYMVYVSQDRLTPSAKH AVAQAITDAHLTHTGEEHS LAQVNFQEQPAGNVFLGG VQQGGDTIFVHGLHREGR SDELKQRLITDIIAKVSIAAD IDPKHIWVYFGEMPASQM VEYGGLGSHHHHHH | 14 |
| T33-22.3A | MDSPIIEANGTLDELTSFIG EAKHYVDEEMKGILEEIQN DIYKIMGEIGSKGKIEGISE DRVAYLLELLLRYEKMVNK SFVLPGGTLESAKLDVCRT IARRAERKVATVLREFGIG KVALRYLKVLERLLFLLARV IEIEKNKEGSHHHHHH | 17.3 |
| T33-22.3B | MSQAIGILELTSIAKGMELG DAMLKSANVDLLVSKTISP GKFLLMLGGDTGAIQQAIE TGTSQAGEMLVDSDLIKDI HPSVLPAISGLNSVDKRQA VGIVETWGVTACIIAADFAV KGSNVTLVRVHMAFGIGG KCYMVVAGDVSDVNNAVD VASRVAGALGLLVYRSVIP RPHEAMWRQMVEGGSHH HHHH | 20.2 |
| T33-24.3A | MPLLKFDLFYGRSDEQIKS LIDAAHGAMVLAFGVPASD RYQTVSQHRPGEMVLEDT GLGYGRTDAVVLLTVISRP RSEEQKVLFNRLLTAALEV LCGISPDDVIVALVENSDA DWSFGGGRAEFLTGDLVG GSHHHHHH | 15.2 |
| T33-24.3B | MSKAKIGIVTVSDRAFAGIY EDISGKAIIDTLNDYLTSEW | 19.9 |

| | |
|---|---|
| | EPIYRVVPDDKDIIVTTLAY MALIEDCCLIVTTGGTGPA KRDVTPEATEAVCDRMMP GFGELMRAESLKFVPTAIL SRQTAGLLGDSLIVNLPGK PKSIRECLDAVFPAIPYCID LMEGPYLECNEAVIKPFRP GSHHHHHH |
